# The mannose 6-phosphate receptor system is ancient, widespread in eukaryotes, and functionally diversified in the human parasite *Trichomonas vaginalis*

**DOI:** 10.64898/2026.10.22.760878

**Authors:** Abhishek Prakash Shinde, Harpreet Kaur, Shih-Yun Guu, Alois Zdrha, Tamara Smutná, Petr Rada, Karel Harant, Kay-Hooi Khoo, Joel Bryan Dacks, Jan Tachezy

## Abstract

In mammals, the mannose 6-phosphate (M6P) pathway tags newly synthesized hydrolases for delivery to the lysosomes. The pathway underlies healthy lysosomal function, but whether an equivalent system operates outside animals remains largely unexplored. Surveying 53 representative species, across eukaryotic supergroups we show that the M6P receptor (MPR), the phosphorylating enzyme GNPT, and the uncovering enzyme (UCE) form a statistically robust, co-occurring module across extant eukaryotic supergroups. In the human parasite *Trichomonas vaginalis*, this module has diversified into an expanded MPR family of seven paralogs, the largest reported within these 53 species, alongside a structurally minimal single GNPT gene, despite the complete absence of UCE. Using co-immunoprecipitation (co-IP), mass spectrometry and confocal microscopy, we demonstrate that MPRs and GNPT localise to the Golgi apparatus, and their interactomes show paralog-specific enrichment of distinct hydrolase cargo, adaptor complexes and the retromer complex proteins for retrieval of MPRs, presenting a full receptor-recycling cycle. Critically, mass spectrometry based glycomics directly detects mature M6P-modified N-glycans in *T. vaginalis* built on unique Man_5_GlcNAc_2_ structure with no trace of GlcNAc-capped intermediate, seen in mammalian cells. Together, these results provide evidence that M6P pathway is an ancient and widespread eukaryotic system, with its functionally diversified variant operating in *T. vaginalis* for lysosomal biogenesis.

## Introduction

Lysosomal biogenesis is essential for the maintenance of eukaryotic cell viability. Lysosomes are the primary degradative and signalling hub of the cell (De Duve, 2005; Yang & Wang, 2021). In humans, their maintenance requires continuous delivery of over 60 soluble hydrolases and over 100 membrane proteins from the endoplasmic reticulum, through the Golgi network and endosomal system, to form a single membrane bound acidic organelle (Braulke & Bonifacino, 2009; Saftig & Klumperman, 2009). Since this route is shared by roughly 2700 different proteins, of which only a small fraction are lysosomal hydrolases, signal-mediated sorting is essential for their selective delivery (Di Lorenzo et al., 2018). In mammals, the mannose 6-phosphate (M6P) signal carried on N-linked glycans of nascent hydrolases, is recognized in the Golgi by transmembrane M6P receptors (MPRs) that divert the cargo away from the default secretory route to the lysosomes (Kornfeld, 1986).

In the mammalian system, generation of the M6P signal is restricted to lysosomal hydrolases that possess an asparagine residue within the consensus N-glycosylation sequon (Asn-X-Ser/Thr), known as the soluble lysosomal targeting sequence (s-LTS). Nascent lysosomal hydrolases containing an N-terminal signal peptide are co-translationally translocated into the ER lumen, where they undergo N-linked glycosylation on the asparagine residue of the s-LTS with a high mannose oligosaccharide (in mammals; Glc_3_Man_9_GlcNAc_2_) (Braulke & Bonifacino, 2009; Ruddock & Molinari, 2006). N-acetylglucosaminyl-1-phosphotransferase (GNPT, encoded by GNPTAB and GNPTG) first transfers a GlcNAc-phosphate group onto the glycan, generating a phosphodiester intermediate (GlcNAc-P-Man). N-acetylglucosamine-1-phosphodiester α N-acetyl-glucosaminidase (NAGPA) also known as the uncovering enzyme (UCE), then catalytically removes the GlcNAc moiety from the phosphodiester intermediate, exposing the mature phosphomonoester M6P ligand (Coutinho et al., 2012; Gorelik et al., 2020; Varki & Kornfeld, 1980). Both phosphodiester and phosphomonoester forms of M6P ligands can be bound to MPRs, of which mammals possess two structurally distinct types; the cation-independent MPR (CI-MPR) carrying 15 mannose receptor homology (MRH) domains and the smaller, dimerizing cation-dependent MPR (CD-MPR) with a single MRH domain (Bohnsack et al., 2009; Coutinho et al., 2012; Höning et al., 1997; Olson et al., 2015). Receptor-cargo complexes are sorted into clathrin-coated vesicles by AP1 and GGA adaptors for direct delivery to endosomes or reach the lysosome indirectly via endocytosis of secreted hydrolases. MPRs are subsequently retrieved from early/recycling endosomes back to the Golgi via the Retromer complex and sorting nexins for further rounds of delivery (Arighi et al., 2004; Seaman, 2004).

Outside mammals, the system has been poorly explored. In 2011, Retromer cargo including CIMPR, GNPTAB and UCE proteins were assessed from a limited sample set of eukaryotes available at the time and showed patchy distribution (Koumandou et al., 2011). Functionally, GNPT and MPR homologs were reported in *Drosophila melanogaster* and *Hydra vulgaris*, which possess MPRs with 5 and 4 repetitive MRH domains, respectively (Bhamidimarri et al., 2018; Hasanagic et al., 2015). While the simplified DmGNPT in *Drosophila* demonstrates phosphotransferase activity, its MPR homolog (LERP) was shown to be implicated in the lysosomal function with no confirmation of a functioning M6P pathway (Dennes et al., 2005; Hasanagic et al., 2015; Li et al., 2024). MPR-like protein (MRL-1) has been reported in yeast, despite no M6P signal on its hydrolases (Whyte & Munro, 2001). In protists, biochemical activity of the GNPT enzyme has been previously reported in *A. castellani*, although no GNPTAB and UCE genes were reported (Ketcham & Kornfeld, 1992). *Dictyostelium discoideum* has been shown to generate distinct, non-canonical phosphorylated glycans including methyl-esterified and sulphated forms not seen in the mammalian M6P system, with the interaction of its modified hydrolases with human CI-MPR (Condie et al., 2025; Marron-Terada et al., 2000). Overall, this leaves uncertainty in the extent to which the mammalian M6P-depedent signalling system is conserved outside of metazoa. Tantalizingly, proteome analysis of phagolysosomes from *Trichomonas vaginalis,* a member of the supergroup Metamonada which is evolutionarily distant supergroup from Metazoa and Amoebozoa, also identified MPR-like proteins (Zimmann et al., 2022). *T. vaginalis* is a human parasite, which causes the most frequent non-viral sexually transmitted disease world-wide. This parasite is highly active in phagocytosis of various cells at vaginal mucosa, including bacteria, erythrocytes, leukocytes, and fragments of damaged epithelial cells. The engulfed cells are rapidly digested in phagolysosomes (José et al., 1998; Midlej & Benchimol, 2010), which are equipped with a large set of cysteine proteases and other lysosomal hydrolases (Zimmann et al., 2022). Consistent with the presence of MPR-like proteins, we previously reported lineage-specific expansion of adaptins and the retromer complexes (Maciejowski et al., 2023; Shinde et al., 2024).

Here, we combine a pan-eukaryotic genomic survey of the M6P pathway across 53 eukaryotes, with an in-depth evolutionary analysis of the parasitic parabasalids, and use co-immunoprecipitation, immunofluorescence microscopy and mass spectrometry to functionally characterize this system in *T. vaginalis*. We show that *T. vaginalis* possesses an expanded set of MPR paralogs alongside a structurally minimal GNPT enzyme that these receptors and enzyme localize to the Golgi compartment, pulldown distinct lysosomal cargo, and associate with the adaptor complex and the retromer complex. Additionally, glycomic analysis directly detects mature M6P-modified N-glycans in *T. vaginalis*. Together, these data reveal a functional M6P signalling system in *T. vaginalis,* widespread presence in diverse eukaryotes and are consistent with the M6P system being present in LECA.

## Results

### A pan-eukaryotic survey reveals a widespread and ancient core M6P system with highly divergent M6P receptors

To examine the extent to which the M6P signalling system exists as a feature of diverse eukaryotes, we conducted a comprehensive genomic survey across 53 eukaryotic species spanning all major supergroups. This sampling was done from the comparative set (TCS) of EukProt v3 database, based on the confirmed presence of AP1 complex (Grech et al., 2025). We assessed the distribution of MPRs, the hexameric GNPT enzyme subunits (GNPTAB and GNPTG), UCE (encoded by NAGPA gene), and the most studied lysosomal hydrolase cargo cathepsin D (Figure 1a). Additionally, we also surveyed the presence of M6P-independent lysosomal hydrolase transporters: Sortilin (VPS10) and LIMP-2 (Extended Data Fig. 1a).

**Figure 1:**
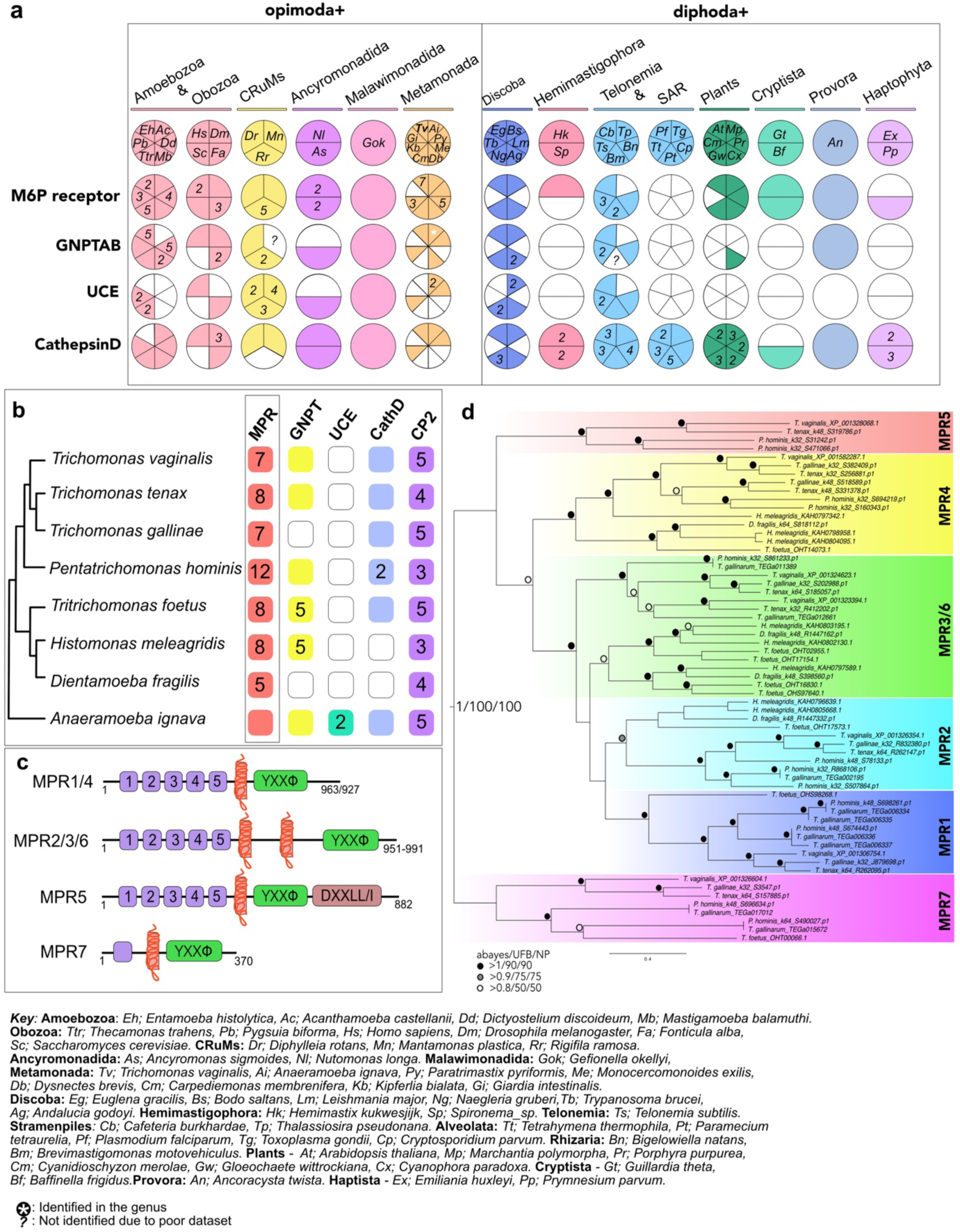
Evolution of Mannose 6-phosphate signalling system. a) Coulson plot representing identification of M6P signalling system protein homologs across selected 53 pan-eukaryotic species, classified with their eukaryotic roots; Opimoda+ and Diphoda+ specific lineages. Colour represents positive identification, blank represents no identification, numbers represent identification of multiple paralogs in a species, star represents identification in a sister species, ‘?’ represents poor incomplete/fragmented dataset. b) Simple plot representing identification of core M6P system and lysosomal enzymes in parasitic parabasalids and free-living Anaeramoeba, colour represents positive identification, blank represents no identification, numbers represent identification of multiple paralogs in a species. MPR homologs are highlighted with a rectangle. c)Variable domain topologies of MPR homologs identified the parabasalid lineage. d) Phylogenetic analysis of parabasalid MPRs, maximum likelihood tree was constructed using IQTree (Best fit model: WAG+F+I+G4) with 62 sequences and 655 positions. The support values provide aBayes posterior probability/ultra-fast bootstrapping/non-parametric bootstrapping. Well supported monophyletic TvMPR clades are represented in shaded colours.

MPR homologs were identified in ∼74% of surveyed eukaryotic species in all the surveyed eukaryotic supergroups. At least one species of all supergroups possesses an MPR homolog. We identified MPRs in all the species of the opimoda+ eukaryotic root (Williamson et al., 2025) such as Amoebozoa, Obazoa, CRuMs, Ancyromonadida and in the single surveyed species of Malawimonadida. While nearly all the species of the Metamonada supergroup retain diverse homologs of MPRs, *Giardia intestinalis* was not found to encode aMPR homolog. Members of the diphoda+ eukaryotic root showed more frequent lack of MPRs, including important parasitic lineages like Trypanosomatida with *Trypanosoma brucei* and *Leishmania major* and Alveolates such as *Plasmodium falciparum*, *Toxoplasma gondii* and *Cryptosporidium parvum*. By contrast, the parasitic parabasalid *T. vaginalis* has the largest MPR complement with seven homologs among the 53 surveyed species.

The GNPTAB gene was confirmed to be present in 26 of the 53 surveyed species (∼49%). GNPTAB followed a similar root-based distribution as MPRs, retained in 17 of 24 opimoda+ species (70%), compared to only 9 of 29 diphoda+ species (31%). To further validate the presence of a putatively functional GNPTAB enzyme, we conducted domain analysis with InterProScan (Table S5) to identify its four stealth domains (CR1-4), with the domains CR1 and CR2 as a requirement for putative active fold formation (Li et al., 2022, 2024). GNPTAB homologs are found in most species of Amoebozoa, Obazoa and CRuMs, except for *A. castellani*, *Saccharomyces cerevisiae* and *Mantamonas plastica,* respectively. Due to the identification of partial GNPTAB sequences, we could not confirm its complete absence in *Mantamonas,* and due to the poor BUSCO score of *Brevimastigomonas motovehiculus*, possibly no GNPTAB homologs were identified in its transcriptome (Grech et al., 2025). By contrast, GNPTG, the non-catalytic subunit of the GNPT hexameric enzyme in mammals, was identified in only 3 of the 53 species, including humans, the stramenopile *Cafeteria burkhardae,* and *Gefionella okellyi* of Malawimonadida. As these latter two identifications could be due to false positive identifications or contamination and may warrant further investigation, the overall lack of GNPTG beyond humans where it has been characterized suggested that it is unlikely a wide-spread facet of the MPR system and it not considered further (Table S1).

The UCE demonstrated a considerably restricted distribution compared to MPR and GNPTAB with only 19 positive species out of 53 (36%), and majority of its occurrence limited to the opimoda+ root. Within the diphoda+ root, only a few discobans such as *Bodo saltans, Naegleria gruberi* and *Andalucia godoyi,* a few members of the Telonemia, Stramenopiles and Rhizaria groups encoded putative homologs. Notably, while *Anaeramoeba ignava* possesses two UCE homologs, none could be found in *T. vaginalis*, even when using the *A. ignava* sequences as queries for homology searches. Despite the reported biochemical evidence of GNPT activity in *A. castellani* (Ketcham & Kornfeld, 1992; Lang et al., 1986), no GNPTAB, GNPTG or UCE encoding genes were identified ((Clarke et al., 2013), this paper), representing a discrepancy between historical biochemical data and genomic evidence. Cathepsin D, a canonical M6P-dependent lysosomal hydrolase (Tavares & daSilva, 2017) was identified across all eukaryotic supergroups while showing opposite root bias to the enzymatic machinery. It is present in 24 of 29 diphoda+ root species (∼83%), including all Alveolata and plant species, compared to which 17 of 24 opimoda+ root species (∼70%), indicating that trafficking of cathepsin D might not depend completely on the M6P-dependent delivery system.

The M6P-independent receptors were more broadly present among diphoda+ species (Extended data Fig. 1a), with sortilin showing relatively higher presence compared to MPRs (19 positive species for sortilin and 18 positive species for MPRs). LIMP2 is sparsely distributed in both diphoda+ compared to opimoda+ species. The reciprocal presence of either the MPR-dependent or MPR-independent complement is most striking at the level of individual parasitic lineages: apicomplexan parasites (*Plasmodium falciparum, Toxoplasma gondii, Cryptosporidium parvum*) retain the M6P independent machinery with sortilin mediated hydrolase sorting, while lacking MPR, GNPT and UCE entirely, whereas metamonads like *T. vaginalis* show the inverse by retaining MPRs while completely lacking sortilin and LIMP2. Along with sortilin, ciliates *T. thermophila* and *Paramecium tetraurelia* also possess LIMP2 for its hydrolase sorting (Extended data Fig. 1a). Interestingly, five of the eight metamonada species nonetheless retained the cargo protein prosaposin and β-glucocerebrosidase reported to be sorted by M6P-independent receptors in humans (Lefrancois et al., 2003; Reczek et al., 2007) (Table S1). The complete comparative genomic searches and the resulting identifications are provided in extended information (Tables S2 and S3).

To test whether MPR, GNPTAB, UCE and cathepsin D function as a coupled system rather than co-occurring by chance, we computed a pairwise 13×13 Jaccard co-occurrence matrix deriving co-occurrence indices across the 53 surveyed species (Extended data Fig.1b) and independently verified both the index values and their statistical significance by Fischer’s exact test on the underlying presence/absence data (Table S1). MPR, GNPTAB and UCE form a significant triad (MPR-GNPTAB: J=0.610, p=0.0015, MPR-UCE: J=0.487, p=0.0005, GNPTAB-UCE: J=0.586, p=0.00003), to be considered as a statistically robust M6P-dependent system module (Extended data Fig. 1b, Table S10). By contrast, while MPR’s co-occurrence with cathepsin D and prosaposin is high, Jaccard indices show poor significance with p=0.24 and p=0.12 respectively suggesting a distribution independent of MPR module. Cathepsin D demonstrates a stronger co-occurrence with sortilin receptor (J=0.6, p=0.017), suggesting that cathepsin D trafficking is statistically linked to both M6P-dependent and M6P-independent routes across the pan-eukaryotic species. Thus, our comparative genomic and co-occurrence analysis has identified the presence of a broadly conserved lysosomal biogenesis system across eukaryotes with clear lineage-specific modifications to be explored.

MPR domain architecture is broadly diversified across eukaryotes. The MRH domain that define MPRs is not restricted to this receptor family, so we required each positive candidate to carry at least one MRH domain, and a tyrosine and/or dileucine cytosolic motif (Castonguay et al., 2011). All 76 MPR sequences identified across the 53 species met these criteria. Additionally, the majority of MPRs demonstrate the presence of at least one transmembrane domain at the C-terminus prior to the tyrosine domain, three homologs, one in each *M. balamuthi, T. trahens* and *D. discoideum* completely lack a transmembrane domain (Extended Data Fig. 2a). Within this diversity of MPRs, *T. vaginalis* stands out with an expanded set of seven paralogs presenting four distinct topologies, each carrying one to five MRH domains, a single transmembrane domain, and a C-terminal tyrosine-based sorting motif (YXXΦ). Additionally, TvMPR5 carries a dileucine motif otherwise restricted to Obazoa as represented by humans and Drosophila along with a rare occurrence in two distinct homologs of *Thecamonas trahens* (Figure 1c, Extended Data Fig. 2a). Meanwhile, across all eukaryotic lineages a common topology of MPRs is observed with a single MRH domain, a transmembrane domain and a following tyrosine sorting motif representing an ancestral MPR topology (Extended Data Fig. 2a).

Due to the repetitive nature of the MRH domains within MPRs and their vast structural diversity, phylogenetic analysis was not a suitable approach to understand their evolutionary relationship. To enrich the evolutionary and statistical analysis, we surveyed the complete TCS dataset of 196 species inclusive of the 53 species shown in Figure 1a, along with all the Parabasalia species sampled (see below), resulting in a total of 329 confirmed MPR homologs. We then conducted CLANS clustering analysis (Extended Data Fig. 3a) and PCA (Principal Component Analysis) supplemented with k-means clustering algorithm (Extended Data Fig. 3b-e). CLANS clustering analysis resulted in three distinct clusters: with single MRH topology MPRs (e.g: human CD-MPRs), multi-MRH topology MPRs (e.g. human CI-MPRs) and parabasalid MPRs forming three distinct clusters (Extended Data Fig. 3a). PCA analysis spatially distributed the 329 MPR sequences (Extended Data Fig. 3b) with the k-means algorithm clustering them into five distinct evolutionary groups (k=5) based on their pairwise sequence similarities with mean similarity distribution of 0.0337 (3.37%) (Extended Data Fig. 3c-e). Cluster 1 with with the human CI-MPR homolog represents the smallest cluster with only 29 sequences, while Cluster 3 with the human CD-MPR homolog represents the largest cluster with 108 sequences, while the parabasalid MPR homologs with five repetitive MRH domains form the Cluster 4 with 66 sequences (Extended Data Fig. 3e, Table S4). Together, this identifies a broadly conserved M6P signalling system across eukaryotes with clear lineage-specific modifications, most strikingly by the parabasalids.

### Deep-dive analysis reveals an expanded set of diverse MPRs in parasitic parabasalids

Because *T. vaginalis* showed an expanded set of MPRs with four domain architectures, and was found previously in a lysosomal proteome, we conducted a more intensively sampled (i.e. deep-dive) comparative genomics survey of M6P-dependent trafficking system in eight parabasalid species and the free-living relative *A. ignava* (Figure1b). This included the survey of core M6P system module (MPR, GNPT and UCE) and lysosomal hydrolases identified in the *T. vaginalis* phagolysosomal proteome previously tested for its lysosomal targeting and localization such as cysteine peptidase (CP2, a cathepsin L), and the conserved cathepsin D (CathD) (Chen et al., 2011; Zimmann et al., 2022). Additionally, we surveyed more lysosomal hydrolases; acid phosphatase (AP), β-hexosaminidase (βHX) and MPR trafficking components; CLN3 and epsinR (Braulke & Bonifacino, 2009; Calcagni’ et al., 2023; Mills et al., 2003; Saint-Pol et al., 2004) (Extended Data Fig. 2b). We found that the complement of MPR genes was expanded in all parabasalids with paralog count ranging from five in *Dientamoeba fragilis* to twelve in *Pentatrichomonas hominis*, with *Trichomonas gallinae* possessing seven paralogs alongside *T. vaginalis*, and *Trichomonas tenax, Tritrichomonas foetus* and *Histomonas meleagridis* each carrying eight paralogs. Meanwhile, the free-living sister *A. ignava* carries only a single ancestral-like MPR homolog. All parabasalid MPRs demonstrate structural homology to the TvMPR topologies shown in Figure 1c, interestingly except for TvMPR5 topology, being the only one to possess a dileucine motif at its C-terminus otherwise restricted to Obazoa. GNPT (synonymous to GNPTAB herein) was detected in most parabasalids except in *T. gallinae* and *D. fragilis*, and UCE was not found in any parasitic parabasalid despite two homologs being identified in *A. ignava*. Cathepsin D was not detected in *H. meleagridis* and *D. fragilis*, with expanded complements of other lysosomal hydrolases AP, CP2 and βHX identified across all the parabasalids ranging from three to five genes each. Trafficking components of MPRs; CLN3 and epsinR demonstrate gene complement expansion comparable to MPRs. CLN3 is present with four and five genes identified in most parabasalids except for single homolog identified in *H. meleagridis* and *A. ignava*. EpsinR showed expanded paralog count ranging from two to five genes in all Tritrichomonadida species and *P. hominis*, with single homolog detected in *T. vaginalis, T. tenax* and *A. ignava*, while undetected in *T. gallinae*.

Because the domain architectures in the parabasalid MPRs were more conserved between taxa, they could be reliably aligned and used for phylogenetic analysis. Maximum likelihood and approximate Bayesian phylogenetic reconstruction resolved the parabasalid MPRs into six well-supported clades with each clade represented by at least one TvMPR, corresponding to different domain-architecture types of TvMPRs (Figure 1c-d). The tree was rooted at the MPR7 clade, exclusively comprising single MRH domain parabasalid MPRs, consistent with the ancestral single-domain topology, also observed with Anaeramoeba MPR homologs (Extended Data Fig. 2a). However, due to high sequence divergence, *A. ignava* MPR homologs were excluded from the analysis. Phylogenetic reconstruction indicates that at least four distinct MPR types (MPR5, MPR2/3/6, MPR4 and MPR7) were present in the last parabasalid common ancestor (LPCA), with subsequent lineage-specific duplications generating the observed expanded repertoire (Figure 1d). To evaluate which of the MRH domains across TvMPR1-7 putatively retain the biochemical capacity for M6P binding, we combined sequence similarity to human MRH domains with conservation of the characteristic QREY tetrad with conserved cysteine residues (>6) and structural alignment supplemented with TM-scores (Extended Data Fig. 5, Table S11). TvMPR5 and TvMPR7 are only predicted to retain binding capacity with moderate confidence, due to the absence of Glutamine from the QREY tetrad, and retention of 6x cysteine residues and TM-scores of 0.28 and 0.59 respectively against their closest human MRH domains (Extended Data Fig. 5). By contrast, the other five MPR paralogs (TvMPR1,2,3,5,6) retain the full QREY tetrad and 6x cysteine residues with TM-align scores ranging from 0.6 to 0.77 against their closest human structural homolog, demonstrating highest similarity to the M6P binding MRH domains of human CI-MPR and CD-MPR capacity.

### TvMPR3 and TvMPR7 localize to the Golgi apparatus and engage distinct lysosomal cargo

Experimental evidence for M6P receptors outside animals is limited, with the most detailed work in Dictyostelium. Studies conducted on *Dictyostelium* previously revealed the presence of M6P-OMe, sortilin is considered as a primary receptor for lysosomal enzyme trafficking, despite it encoding MPR homologs. Lysosomal hydrolases were shown to interact with human CI-MPR in this model protist, suggesting the possibility of a MPR receptor sorting system (Condie et al., 2025; Marron-Terada et al., 2000). Given the presence of key components of M6P system in *T. vaginalis*, and yet the absence of some machinery considered essential such as the UCE, we proceeded to validate and characterize the MPR machinery in this model system. For the experimental validation we selected TvMPR3 and TvMPR7, representing structurally distinct paralogs (five MRH domains versus one). Both receptors were expressed with 2xHA and 2xV5 tag in *T. vaginalis* TV17-48 strain to test their localization, with use of 2xV5 variants in case of co-localization with a 2xHA marker (Figure 2). In mammalian cells, MPRs are observed within the post-Golgi trafficking including the TGN, early endosomes and limited distribution on the surface. The parabasalid lineage is defined by the Golgi apparatus associated with the parabasal filaments. An in-house developed antibody against protein disulfide isomerase (α-PDI) labelled the Golgi body (G) associated with parabasal filaments (Pf) forming a semi-V-shaped eight like structure in proximity of the parabasal filament (Extended Data Fig. 4a). Therefore, we used α-PDI labelling as a Golgi marker in *T. vaginalis* cell. Strikingly, both TvMPR3 and TvMPR7 (Extended Data Fig. 4b) co-localized with PDI at the Golgi body, with additional punctate distribution throughout the cell (Figure 2a-b).

**Figure 2:**
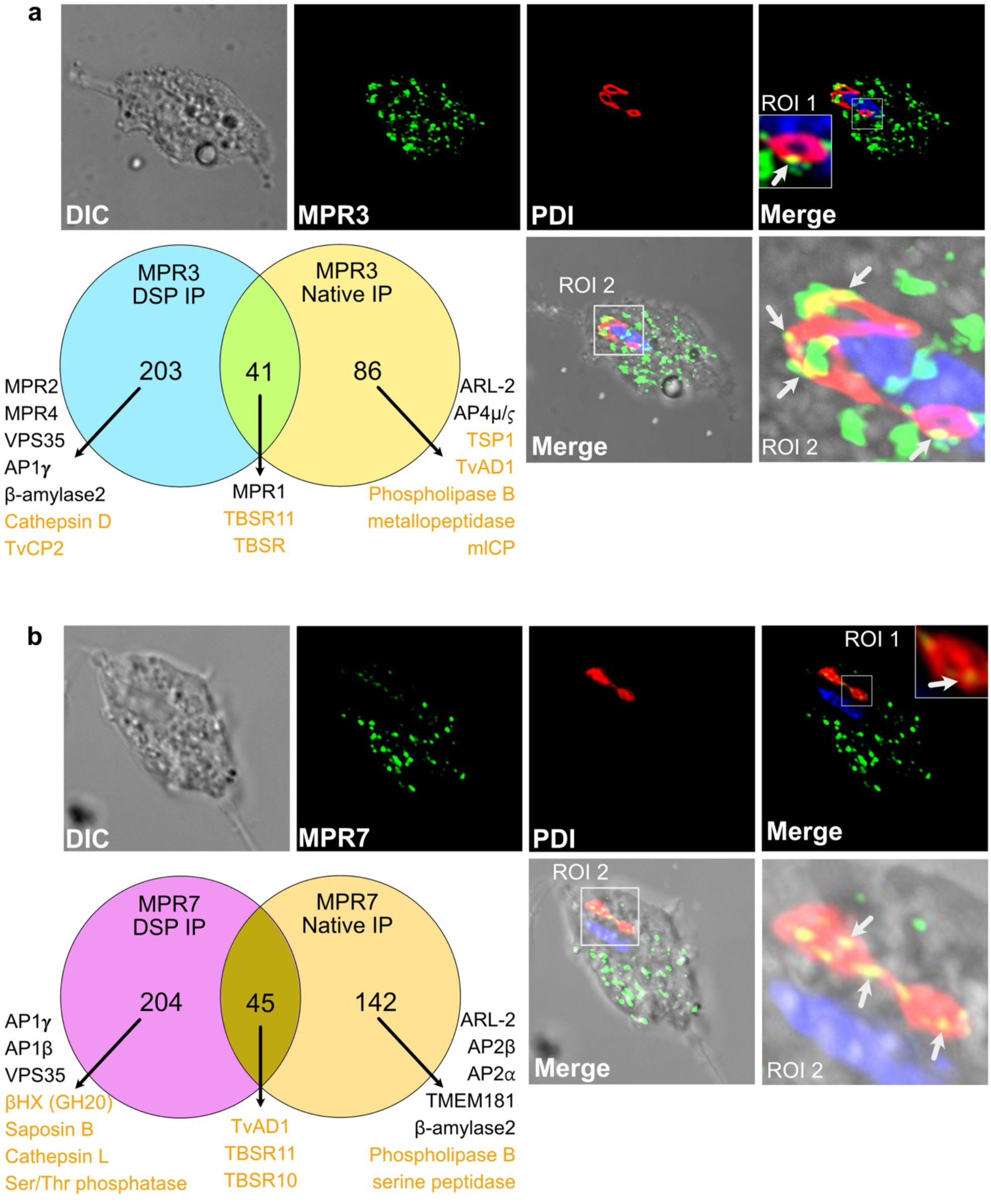
Immunofluorescence microscopy and immunoprecipitation of TvMPRs. a) Confocal immunofluorescence microscopy of 2xHA-tagged TvMPR3 (green channel) with PDI Golgi marker (red channel). Merged channels show overlapping signals in *T. vaginalis* cell with ROI 1, showing a zoomed Golgi sub-domain labelled by PDI, and ROI 2 showing the complete Golgi labelled by PDI. Arrows point co-localizing regions for both proteins. Venn diagram represents LFQ MS identifications by native and DSP cross-linked co-IP methods; number of identifications by each method are provided with significant protein named; phagolysosomal proteins are labelled in orange text. b) Confocal immunofluorescence microscopy of 2xHA-tagged TvMPR7 (green channel) with PDI (red channel). Merged channels show overlapping signals in *T. vaginalis* cell with ROI 1 showing a zoomed Golgi sub-domain labelled by PDI, and ROI 2 showing the complete Golgi labelled by PDI, arrows represent co-localizing regions for both proteins. Venn diagram represents LFQ MS identifications by native and DSP cross-linked co-IP methods, number of identifications by each method are provided with significant detections named, phagolysosomal proteins are labelled in orange text.

Co-immunoprecipitation (co-IP) coupled with label-free quantitative mass spectrometry (LFQ-MS) for both the TvMPRs, performed under both native and DSP cross-linked conditions, identified overlapping and distinct interactions (Figure 4a-b). TvMPR3 co-IP detected 86 significantly enriched proteins unique to native conditions and 203 unique to DSP cross-linking, with 41 overlapping interactions (Figure 4a). TvMPR7 co-IP identified 142 enriched proteins under native conditions and 204 under DSP cross-linking, with 45 overlapping interactions (Figure 4b). Volcano plots demonstrating enriched proteome is provided (Extended Data Fig. 6) with full quantitative dataset for both receptors (Table S6-7). Both the receptors demonstrate interactions with lysosomal hydrolases and proteins previously identified (Zimmann et al., 2022). While TvMPR3 uniquely interacts with cathepsin D and metacaspase-like cysteine peptidase (mlCP), TvMPR7 interacts with βHX (Glycosylhydrolase 20/GH20) and cathepsin L. Both receptors interact with lysosomal cargo such as Trichomonas beta sandwich repeat 11(TBSR11) and secreted protein β-amylase 2 (Stáfková et al., 2018). Interestingly, TvMPR3 interacts with other MPRs like TvMPR1,2 and 4. Notably, both TvMPR3 and TvMPR7 showed interactions with distinct paralogs of AP1/AP2 Adaptor complex proteins responsible for its anterograde trafficking, and the Retromer complex proteins (VPS35 and VPS26A/B) for retrograde recycling of the MPRs.

### TvMPRs interact and co-localize with the Retromer complex proteins

For the receptor-mediated sorting cycle to be functionally complete, receptors are required to be retrieved from the endomembrane compartments to be returned to the *trans*-Golgi network for reuse rather than degradation in the acidic lysosomes after a single round of cargo delivery. We therefore tested co-localization of TvMPR3 and TvMPR7 with the previously tested Retromer complex subunit VPS26A/B (Shinde et al., 2024) by confocal immunofluorescence microscopy. VPS26A/B showed strong co-localization with both, TvMPR3 (Pearson’s R = 0.7, Manders’ M1 = 0.995, M2 = 0.993) and TvMPR7 (Pearson’s R = 0.611, Manders’ M1 = 1.0, M2 = 0.992) (Figure 3a-b). The presence of VPS26A/B and VPS35 in TvMPR3 co-IP and VPS35 in TvMPR7 co-IP datasets (Extended Data Fig. 6a-b), provide proteomic and microscopic evidence for Retromer engagement.

**Figure 3:**
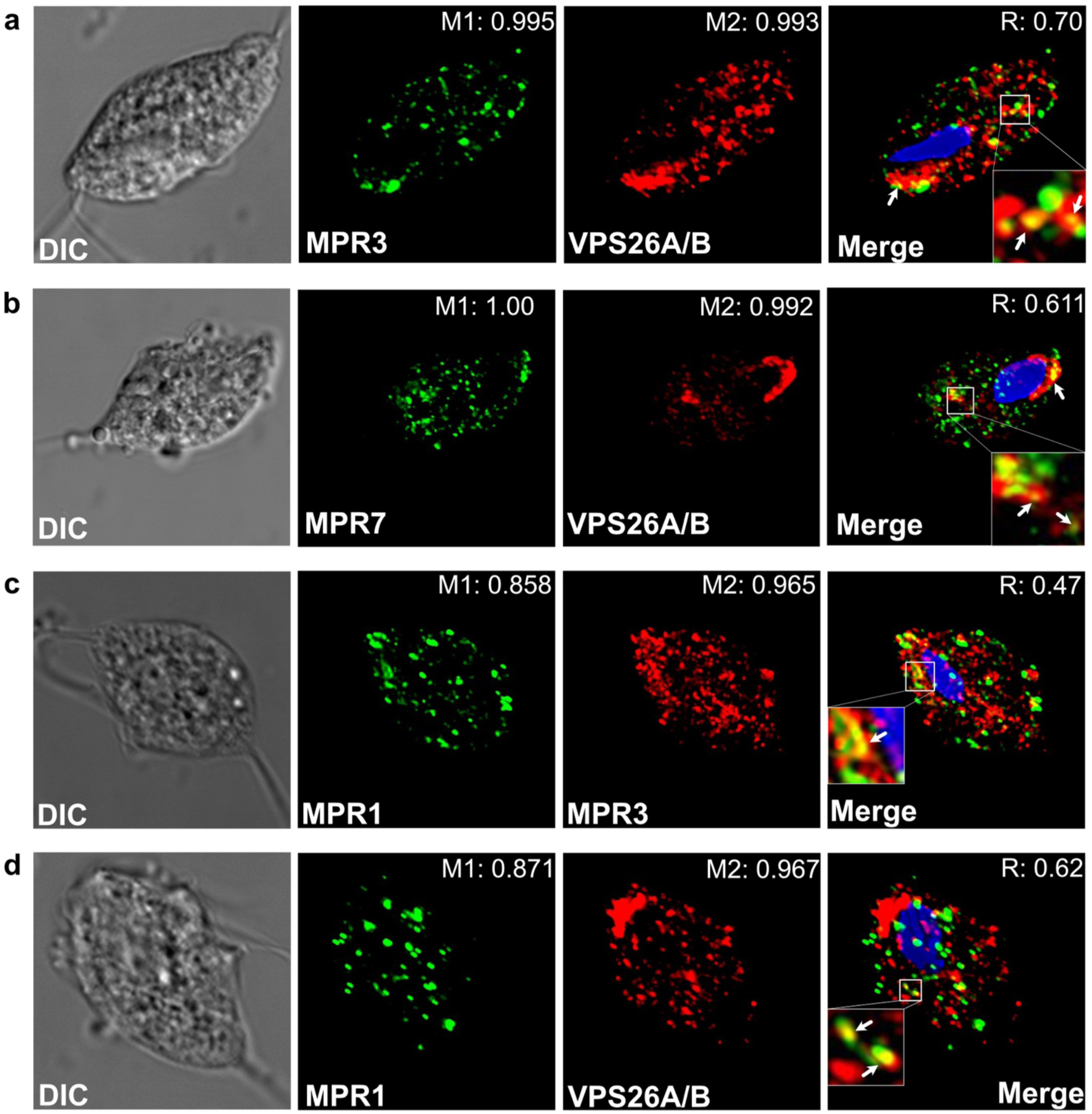
Co-localization analysis of TvMPRs with the Retromer complex protein VPS26A/B. a) Confocal immunofluorescence microscopy for co-localization of 2xV5 tagged TvMPR3 (green channel) with 2xHA tagged VPS26A/B (red channel) of the Retromer complex labelling the recycling endosomes and Golgi subdomains in *T. vaginalis* (Shinde et al., 2024). b) Co-localization of 2xV5 tagged TvMPR7 (green channel) with 2xHA tagged VPS26A/B (red channel) c) Co-localization of TvMPR1 (labelled with α-TvMPR1 antibody – green channel) with 2xHA tagged TvMPR3 (red channel) d) Co-localization of TvMPR1 (green channel) with 2xHA tagged VPS26A/B (red channel). Merged channels show overlapping signals in the complete cell, white square represents region of interest with zoomed ROI and arrows demonstrating co-localization, voxel-analyses generating values supporting co-localization of both channels are presented as Pearson’s co-efficient (R), Mander’s co-efficients (M1 and M2) in both the panels.

To determine whether this reflects a general property of TvMPR family rather than feature specific to TvMPR3 and TvMPR7, we examined TvMPR1, a third, structurally distinct paralog also identified in the co-IP dataset of TvMPR3 under both native and cross-linking conditions (Figure 2a). NativeTvMPR1 labelled with in-house anti-MPR1 antibody co-localized with both TvMPR3 (Pearson’s R = 0.47, Manders’ M1 = 0.858, M2 = 0.965) and VPS26A/B (Pearson’s R = 0.62, Manders’ M1 = 0.871, M2 = 0.967) (Figure 3c-d). Based on the findings, we speculate that our data are consistent with the idea that TvMPR1 shares the Golgi and vesicular compartment subdomains with TvMPR3 and that Retromer-mediated recycling is a shared, conserved feature across structurally divergent TvMPR paralogs, functionally completing the receptor trafficking cycle in *T. vaginalis* in a homologous manner to Retromer-dependent MPR retrieval in mammals.

### TvGNPT (GNPTAB homolog) retains the phosphotransferase catalytic fold and localizes in the Golgi apparatus

Having established that the receptor side of the M6P system is extensively diversified and functional, we examined the enzyme generating the M6P signal itself (Figure 4). Our comparative genomics survey identified a single GNPTAB homologous gene in *T. vaginalis* to be referred to as TvGNPT herein (Figure 1). The human GNPTAB precursor (1,256 aa) carries two flanking transmembrane domains and four conserved catalytic regions (CR1-4), along with four additional mammalian-specific regulatory domains (Gorelik et al., 2022). At ∼half the length, the *D. Melanogaster* GNPTAB homolog (DmGNPT, 666 aa) still retains the four CR regions and both the transmembrane domains (Du et al., 2022). By contrast, TvGNPT (343 aa) retains only a single N-terminal transmembrane domain with CR1 and CR2 of the core catalytic region (Figure 4b), placing it in a small group of 35 CR1-CR2-only sequences identified genome-wide in our broader classification of 368 identified homologs in the 196 species of TCS (Table S5).

**Figure 4:**
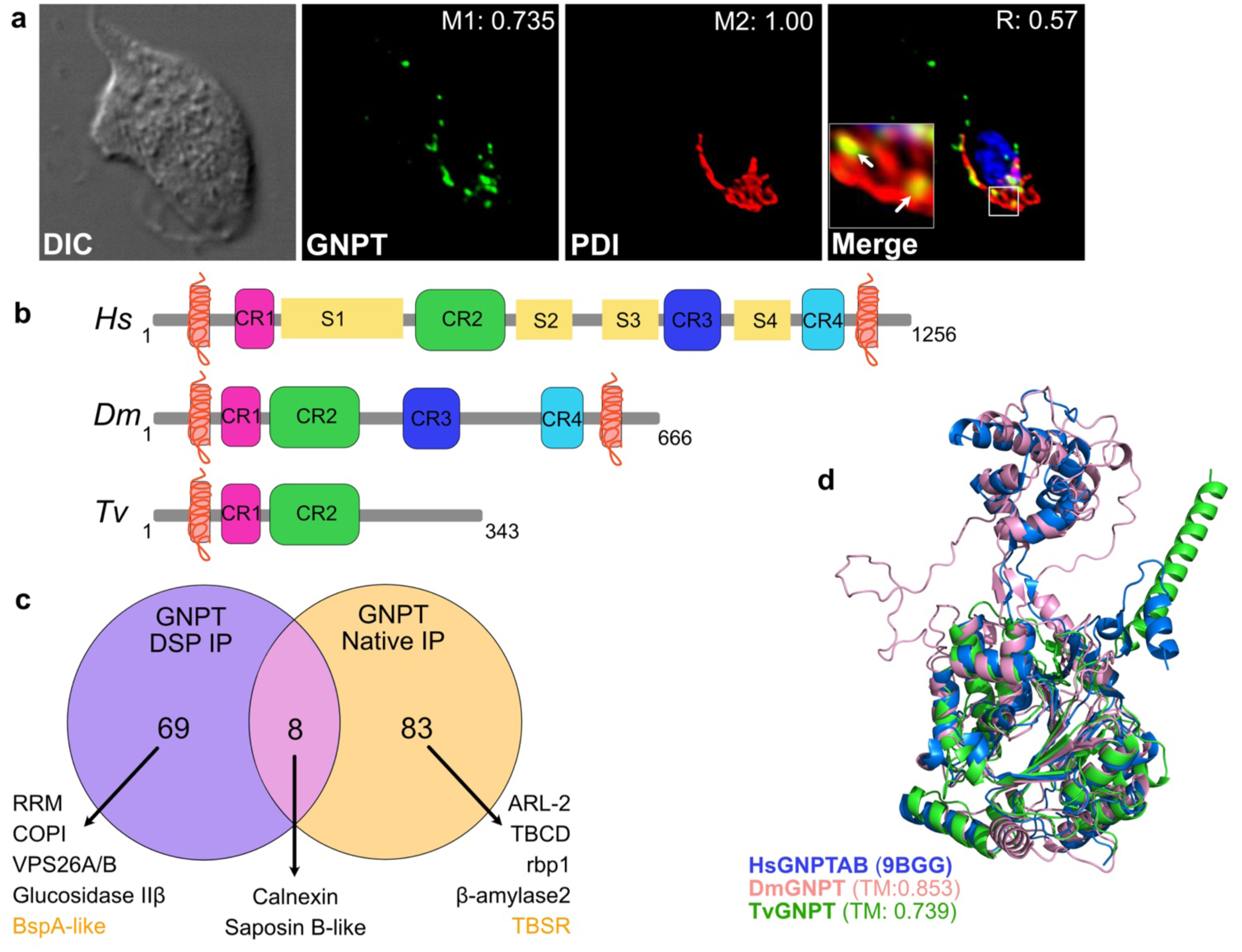
Functional characterisation of the compact TvGNPT enzyme. a) Confocal immunofluorescence microscopy of 2xHA tagged TvGNPT (green channel) and its co-localization analysis with PDI (labelled with α-PDI antibody - red channel). Merge channel shows overlapping signals in the complete cell, white square represents region of interest with zoomed ROI and arrows demonstrating co-localizing Golgi domains and co-localization analysis generating values supporting co-localization of both channels are presented as Pearson’s co-efficient (R), Mander’s co-efficients (M1 and M2) in both the panels b) InterProScan generated graphical demonstration of comparative domain conservation in Human, Drosophila and *T. vaginalis* GNPT enzymes c) Immunoprecipitation experiment Venn diagram represents LFQ MS identifications by native and DSP cross-linked co-IP methods, number of identifications by each method are provided with significant identifications, phagolysosomal proteins are labelled in orange text d) 3D structural alignment of predicted *T. vaginalis* GNPT enzyme and *D. melanogastor* GNPT enzyme with crystal structure of Human active GNPT domain (PDB: 9BGG). Mathematical validation of structural alignments was calculated with TM-align represented with TM-score in reference to HsGNPTAB.

Despite this compact architecture, AlphaFold3-predicted structures of TvGNPT and DmGNPT aligned to the crystal structure of the active human GNPTAB catalytic domain (PDB: 9BGG, (Li et al., 2024)) with TM-align score of 0.739 and 0.853 respectively, both well above the 0.5 threshold for conserved fold (Figure 4d), indicating that the phosphotransferase catalytic architecture is structurally conserved despite the absence of mammalian-specific regulatory domains such as RNA-recognition motifs (RRM1-2) and DNA-methyltransferase associated 1 (DMAP-1) binding domain within the S1-4 regions (Gorelik et al., 2022). While CR3-4 are essential for the formation of catalytic fold, their function is to maintain the structural integrity. Despite TvGNPT missing the CR3-4 regions maintains the catalytic fold structure. In mammalian systems, GNPT is found primarily within the Golgi body. Recombinantly tagged and expressed TvGNPT localized to the Golgi apparatus, marked by anti PDI Ab (Pearson’s R = 0.57, M1=0.735, M2=1.00), along with additional vesicular presence (Figure 4a). This distribution of TvGNPT parallels with the subcellular presence of human GNPTAB (Li et al., 2022, 2024).

Co-IP LFQ-MS of TvGNPT identified 83 proteins significantly enriched under native conditions and 69 under DSP cross-linking, with 8 overlapping identifications (Figure 4c, Extended Data Fig. 6, Table S8). Native co-IP recovered RNA binding protein (RBP-1) possessing two RRM domains, and TIP20 homolog (a COPI-associated Golgi-ER recycling factor), phagolysosomal TBSR and the secreted β-amylase 2, while DSP cross-linking additionally captured interactions with another RRM domain protein, COPI coat complex proteins, glucosidase IIβ (a MRH domain containing protein), calnexin, phagolysosomal BspA-like protein and the VPS26A/B of the Retromer complex, consistent with interactions observed with the TvMPRs.

### Integrated interactome mapping links the TvGNPT and TvMPR trafficking networks

Combining the co-IP datasets for TvGNPT, TvMPR3 and TvMPR7 into a single classified interactome network (Figure 5a,Table S9) resolved 706 proteins across all baits and conditions with 117 TvGNPT-specific, 187 TvMPR3-specific and 246 TvMPR7-specific, 113 shared between both TvMPRs, 19 shared among all three baits, and smaller sets shared pairwise between TvGNPT and each TvMPR (11 with TvMPR3 and 13 with TvMPR7) (Extended Data Fig. 9a).This network included 198 phagolysosomal proteins and 23 proteins previously shown to be dynamically secreted by *T. vaginalis* (Stáfková et al., 2018; Zimmann et al., 2022) (Extended Data Fig. 9b-c), indicating that the M6P signalling system is involved in both the conventional secretory pathway and unconventional, phagolysosome-mediated secretory pathway previously described in this parasite (Zimmann et al., 2022).

**Figure 5:**
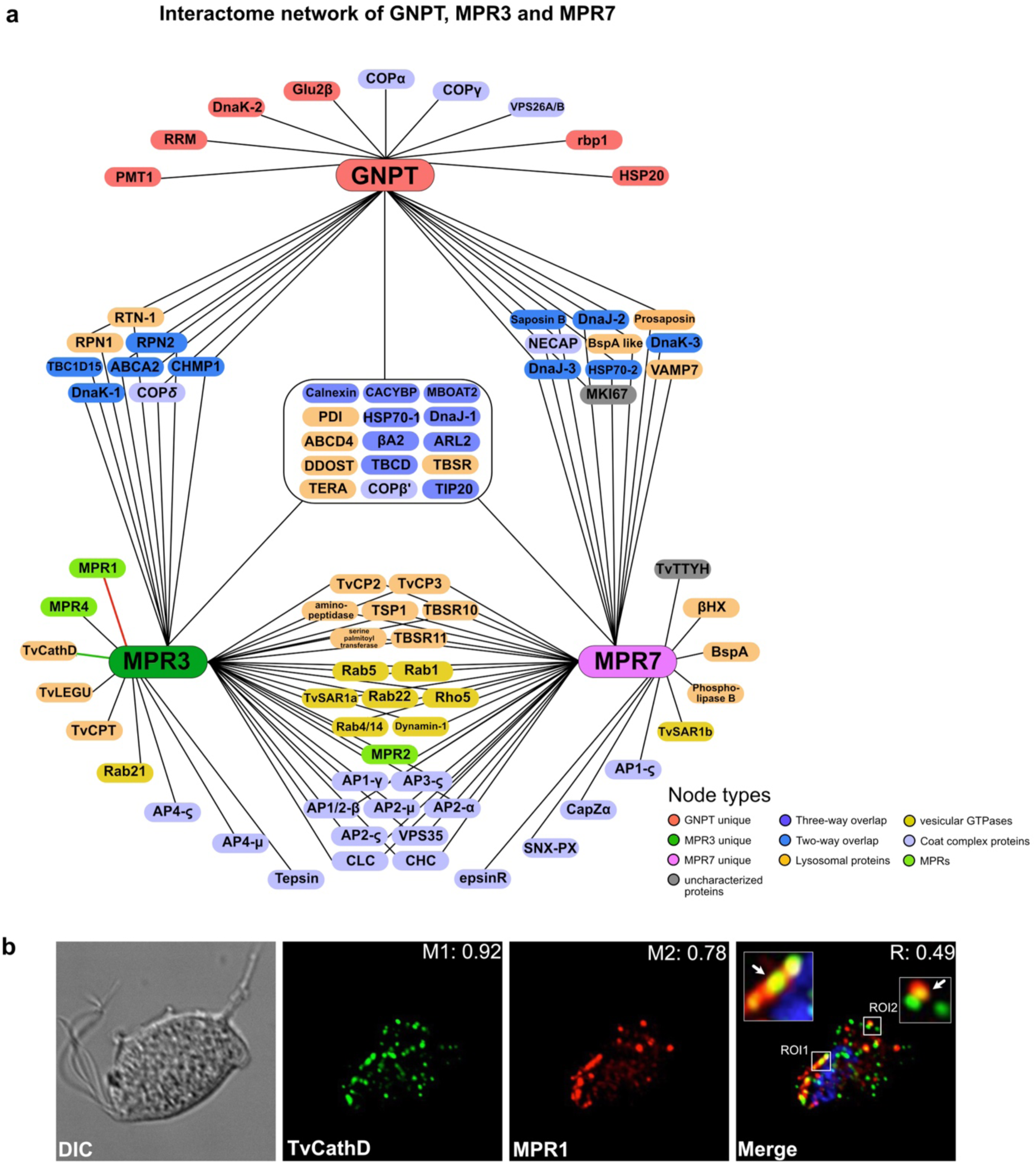
Interactions of the M6P signalling system in *T. vaginalis.* a) Interactome network of TvGNPT, TvMPR3 and TvMPR7. The interactome was manually constructed based on coIP and mass spectrometry experiments, large, coloured boxes indicate baits that were used for the coIP. Unique and shared interactions with the respective baits are shown in solid lines. Interacting partners are colour coded to match their putative functions, and their bait overlap, lysosomal proteins are coloured in orange, uncharacterised proteins are coloured in grey. All interactions are replicated from Extended data Table S9; b) Microscopic validation of shared localisation of most upregulated MPR3 interacting partners, TvMPR1 (red channel) and 2xHA tagged CathepsinD (TvCathD-green channel), its co-localization analysis generating values are presented as Pearson’s co-efficient (R), Mander’s co-efficients (M1 and M2). ROI 1 represents co-localization in the Golgi domain and ROI2 represents co-localization in the peripheral vesicles, overlapping signals are marked with arrows.

Within this network, TvGNPT uniquely interacts with RNA-binding proteins RRM and RBP-1, Glucosidase IIβ, COPI trafficking proteins COPα and COPγ while sharing interactions of lysosomal cargo such as TBSR, prosaposin and BspA-like protein with both TvMPRs. Interestingly, TvGNPT and TvMPR3 interact with a Ribophorin 1 homolog, a subunit of the OST complex involved in N-linked glycosylation and the lysosomal biogenesis localized in the human ER and lysosomes, also identified in *T. vaginalis* phagolysosome (Basrur et al., 2003; Chi et al., 2006; Zimmann et al., 2022) along with Ribophorin 2 homolog of the ER. Additionally, the overlap also includes ATP-binding cassette superfamily proteins of the lysosomes such as ABCD4 and ABCA2 (ATP-binding cassette sub-family A) homologs, other OST complex proteins like DDOST, COPI coat proteins such as COPδ, COPβ’ and its accessory component TIP20 involved in Golgi to ER retrograde trafficking (Kraynack et al., 2005). Notably, the conventionally secreted β-amylase 2, previously shown to demonstrate N-linked glycosylation (Stáfková et al., 2018) is observed to interact with TvGNPT and both the TvMPRs suggesting a primary role of the M6P-signaling system in conventional secretory pathway.

The enrichment of lysosomal hydrolases, resident and secreted proteins in the interactome network such as TvCP2, BspA-like, phospholipase B, TBSR and TBSRs 5, 9, 10 and 11 despite its bait-specificity (Figure 5a, Extended Data Fig. 9b, Table S9) suggest a role of M6P-dependent signalling system in unconventional protein secretion via phagolysosomes. Curiously, the three-way interactome revealed association of M6P system components with a single paralog of *T. vaginalis* putative PDI (TVAG_117470) (Figure 5a, Table S9), amongst three putative PDI paralogs previously found to be enriched in its phagolysosomal proteome with predicted sites for N-linked glycosylation (Zimmann et al., 2022), suggesting a possible role of *T. vaginalis* M6P system in ER-phagy (Sun et al., 2023). While both TvMPR3 and TvMPR7 interact with lysosomal hydrolases such as TvCP3, an aminopeptidase, TBSRs 10-11, and TSP1, TvMPR3 additionally shows exclusive interaction with cathepsin D, and other TvMPRs (TvMPR1,4), not shared with TvMPR7. To determine whether this TvMPR3-cathepsin D interaction reflects a genuine, paralog-specific trafficking route, we co-localized recombinantly tagged cathepsin D with endogenous TvMPR1 (of TvMPR3 specific interactions) using immunofluorescence microscopy (Figure 5b). Cathepsin D showed localisation to the Golgi compartment sharing subdomains with TvMPR1, along with vesicular co-localization with TvMPR1 in the periphery of the cell (Pearson’s R=0.49, M1=0.92, M2=0.78). The complete interactome coupled with confocal immunofluorescence imaging positions TvGNPT and TvMPRs of the M6P signalling system within an interconnected post-Golgi trafficking network for lysosomal biogenesis and protein secretion in *T. vaginalis*.

### Glycomic profiling provides direct evidence for M6P signal formation in *T. vaginalis*

The above data shows that the MPR machinery localizes to the correct locations and interacts with the lysosomal hydrolases and its trafficking proteins to support functional homology with metazoan M6P systems. However, the absence of the UCE from every parabasalid genome (Figure1b) still raises questions. To directly test whether *T. vaginalis* generates M6P-modified N-glycans, we initially screened the PNGase F-released N-glycans from *T. vaginalis* lysate by MALDI-MS in negative mode after permethylation, which is optimal for detection of acidic phosphate group. This analysis identified a series of N-glycans with glycosyl compositions consistent with carrying an additional M6P modification. Subsequent LC-MS/MS analysis unambiguously confirmed the presence of a terminal P-6Hex-carried on a common Man_5_GlcNAc_2_ core structure, which can be further substituted with an additional Pent, HexNAc, or Hex-HexNAc residues (Figure 6, Extended Data Fig. 8). Compared with the M6P-carrying N-glycans typically identified in mammalian samples by the same LC-MS/MS glycomic analysis (Extended Data Fig. 7), it is clear that the underlying Man_5_GlcNAc_2_ structure is unique to *T. vaginalis* due to its lack of further mannosyl extension on the 6-arm of the trimannosyl core in the biosynthesis of its dolichol-linked precursor (Samuelson et al., 2005). Consequently, for the Man_5_GlcNAc_2_ structure with a M6P-modified linear P6Man-2Man on the 3-arm, additional GlcNAc (and Gal-GlcNAc) can only be extended from its 6-arm to generate the so-called pseudo-hybrid type structures reported previously (Paschinger et al., 2012). Intriguingly, MS^2^ on all major glycan peaks detected in negative mode failed to identify any UCE-unprocessed GlcNAc-P-modified Man_5_GlcNAc_2_ and its related structures (Extended Data Fig. 8), whereas a GlcNAc-P-Man_6_GlcNAc_2_ could be readily detected in mammalian cells (Extended Data Fig. 7). Since uncovering this intermediate is the dedicated function of UCE, an enzyme entirely missing from the parabasalid genomes, this result indicates that *T. vaginalis* completes this step so efficiently that the phosphodiester intermediate does not accumulate in detectable amounts, implicating an as yet unidentified enzyme in this step of the M6P pathway. Combined with the Golgi localization and receptor-cargo interactions already established by coIP, this glycomic evidence gives direct chemical proof for M6P mediated lysosomal sorting in this parasite.

**Figure 6:**
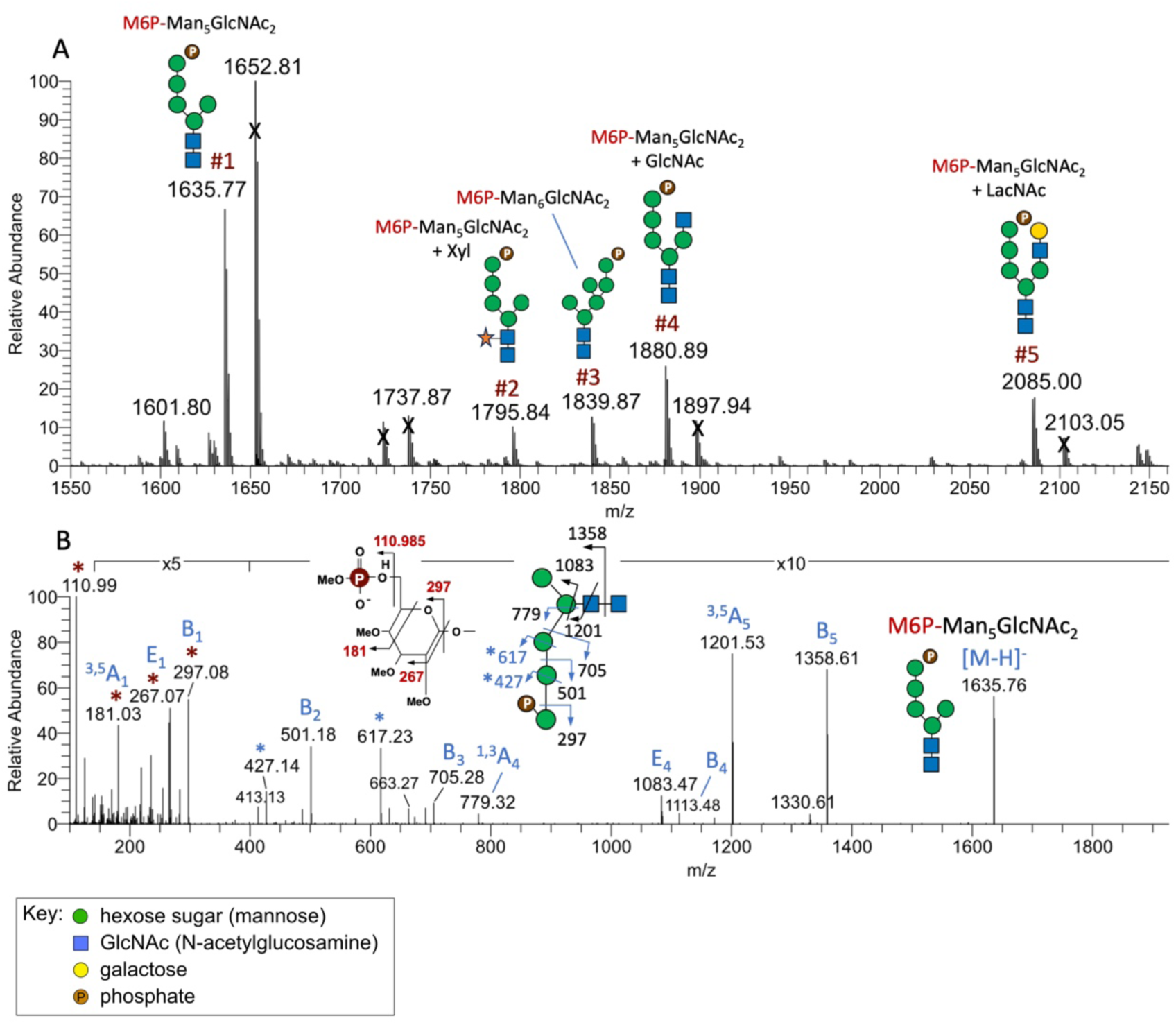
Identification of M6P-carrying N-glycans from *T. vaginalis* by LC-MS/MS analysis. a) MS scans averaged across the entire time range in which the putative permethylated M6P-carrying N-glycans eluted revealed 5 major peaks. Their structures and location of M6P were established by manual interpretation of their respective MS^2^ spectra in comparison with those of similarly processed M6P-carrying N-glycans from CHO cells (Extended data Fig. 8). Fragmentation patterns were illustrated by cartoon drawings using the standard symbol nomenclature (Varki et al. 2015), and the assigned MS^2^ ions were annotated accordingly. Peaks marked with X in (A) did not afford any of the diagnostic MS^2^ ions for the M6P or GlcNAc-P. b) The fully annotated MS^2^ spectrum for the most abundant M6P-Man_5_GlcNAc_2_ component (peak #1) is shown here, while those of peaks #2-5 are shown in Extended data Fig. 7.

## Discussion

The discovery of lysosomes by Christian de Duve over seventy years ago initiated a field that has since transformed our understanding of eukaryotic cell biology. Yet, despite extensive characterization of the M6P-dependent signalling system in mammals, equivalent functional analyses outside of metazoa have been scarce. The most complete characterization has been in *Dictyostelium* which showed both similarity and difference to the mammalian system (Condie et al., 2025). Previously, our identification of MPR-like proteins in the *T. vaginalis* phagolysosomal proteome suggested a possible M6P-dependent lysosomal biogenesis (Zimmann et al., 2022). This work presents a complete evolutionary reconstruction of M6P-dependent and M6P-independent signalling systems, along with the functional proof of the M6P-dependent signalling system in *T. vaginalis*.

Our pan-eukaryotic survey shows that the M6P machinery and its principal cargo are broadly retained across eukaryotic diversity, indicating its presence in LECA, though the precise root-level distribution should be interpreted with caution given potential sampling bias or genome completeness across the surveyed species. The clearest distinction of root-level distribution is the reciprocal retention pattern in two major parasitic lineages; apicomplexan parasites of the diphoda+ retain the M6P-independent machinery while lacking the M6P system, and metamonad parasites, including *T. vaginalis* of opimoda+ show the inverse pattern. The lysosomal hydrolase cargo; Cathepsin D is retained in both apicomplexan parasites and metamonad species, regardless of the receptor for its sorting. This indicates that evolution of the hydrolase receptor system and the hydrolase cargo it delivers are independent events, suggesting that LECA presumably possessed both M6P-dependent and M6P-independent modes of sorting lysosomal hydrolases.

The Jaccard co-occurrence test coupled with Fischer’s exact test (Figure S1B) confirms that MPR, GNPTAB and UCE form a genuinely coupled module (all p < 0.002). However, while MPRs highly co-occur with cathepsin D and prosaposin, the significance of their exclusivity is relatively poor (p=0.24 and p=0.12), supplementing evidence of independent evolution rather than functional coupling to MPRs, clearly observed by reciprocal retention patterns in apicomplexan and metamonad parasites. The loss of one mechanism is generally mitigated by the presence of another mechanism in all the eukaryotic groups, as can be observed in Extended Data Fig. 1. While most eukaryotes sampled showed the presence of at least one type of receptor for lysosomal sorting, some exceptions such as the parasitic protists *G. intestinalis, T. brucei* and *L. major* did not retain any receptor suggesting a possibly distinct pathway. However, interestingly, the retromer complex has been widely demonstrated to be conserved across all these eukaryotes including *Giardia* and Trypanosomatids, despite the absence of a clear receptor (Koumandou et al., 2011).

The comparative genomic picture that emerges in *T. vaginalis* is a paradox pairing a single paralog of TvGNPT retaining only CR1-CR2 catalytic fold, with the most expanded set of MPR paralogs across eukaryotes. The seven MPR paralogs are classified into four ancestral structurally distinct homologs found in the last parabasalid common ancestor (LPCA). Cargo specificity in this system therefore appears to occur downstream of the M6P signal generation step, through distinct complement of MRH domains, sorting motifs and adaptor interactions carried by each receptor paralog rather than the M6P signal formation. TvMPR3 and TvMPR7 illustrate this directly, despite sharing retromer-mediated recycling, they engage distinct lysosomal hydrolase cargo, distinct AP1/AP2 paralogs and distinct adaptor accessory factors Tepsin and epsinR respectively (Figure 5a, Table S11), demonstrating paralog level functional specialization in *T. vaginalis*. That different paralogs of the MPR system bind distinct paralogs of adaptin complexes gives the first hint at a functional explanation to the extraordinarily expanded adaptin complex complement in *Trichomonas* (Maciejowski et al., 2023) and raises the possibility of cryptic diversity of post-Golgi trafficking networks in this parasite.

The integrated interactome extends this picture beyond the core M6P system, its cargo and trafficking components. The detection of Ribophorin 1 and Ribophorin 2 of OST complex involved in N-linked glycosylation, additional OST subunit; DDOST, COPI coat components, and ABC transporters of the lysosome (ABCD4 and ABCA2), together in association of the secreted, N-glycosylated β-amylase 2, suggest that the M6P system in *T. vaginalis* is not an isolated lysosomal targeting module but is embedded within a broader secretory and glycosylation network. The overlap of 198 phagolysosomal proteins and 23 dynamically secreted proteins in this interactome (Figure S6) further suggest a role for M6P-signaling system in both conventional and unconventional (phagolysosomal mediated) secretion in this parasite, a connection previously unexplored or described is relevant to how *T. vaginalis* delivers hydrolytic and potentially virulence-associated proteins to its extracellular environment during infection.

The glycomic data provides direct evidence for the presence of M6P modification on N-glycans. The unusual structure of *T. vaginalis* glycans (Man_5_GlcNAc_2_) reflects a reduced machinery catalysing biosythesis of dolichylpyrophosphate-anchored precursor at ER membrane (Samuelson et al. 2005; Paschinger et al., 2012). While in most eukaryotes, Man_5_GlcNAc_2_, is formed at the cytosolic face of ER, by the asparagine-linked glycosylation (ALG) enzymes, and then flipped to ER lumen where the precursor is extended and branched, the luminal ALGs are absent in *T. vaginalis* (Samuelson et al., 2005, Andrew et al., 2026). Consequently, Man_5_GlcNAc_2_ represent the major oligosaccharide, which is transferred to nascent proteins in ER and available for further modification with pentoses, phosphoethanolamine, and putative phosphate (Paschinger et al., 2012, this study). However, the site of the phosphorylation has not been studied, and particularly the exact position of phosphate modification was not resolved until our analysis. Cellular localization of TvGNPT, and partial colocalization with MPRs indicate that M6P modification machinery is conserved in Golgi compartment structures as in metazoan. Enrichment of phosphorylated glycans and their MS/MS analysis identified the diagnostic M^2^ ion (M6P) Man-2Man-2Man-moiety. Interestingly, all M6P-carrying glycan we detected in *T. vaginalis* was found in its uncovered mature phosphomonoester form, with no evidence of intermediate M6P-GlcNAc equivalent to mammalian phosphodiester tag. Since UCE is not detectable in *T. vaginalis* genome even using the UCE protein sequence from its free-living sister *A. ignava* as search query, this result cannot be explained by a simple retention of a divergent UCE ortholog. Instead, it suggests that an unidentified enzyme catalyses an efficient uncovering of a mature M6P signal. Identifying this enzyme, potentially among the uncharacterized proteins in the TvGNPT and TvMPR interactome, will be an important direction for future work. The underlying glycan itself is built on a Man_5_GlcNAc_2_ core distinct from the mammalian default scaffold, consistent with the parasite’s previously reported, evolutionary reduced N-glycan biosynthesis machinery (Paschinger et al., 2012; Samuelson et al., 2005).

In conclusion, our comprehensive study supports the idea that the M6P system acts in diverse eukaryotes and was present in the last common ancestor of extant eukaryotic species. Different lineages have since then preserved, lost or functionally rebuilt the system for adaptation. In the globally impactful parasite, *T. vaginalis* this manifests as a functional diversification of TvMPRs complements, the presence of a compact TvGNPT enzyme, and the absence of a standard UCE enzyme. This paradigm reveals MPR trafficking routes via distinct adaptor complexes, and conservation of retromer-mediated receptor recycling in *T. vaginalis* (Extended Data Fig. 9d) and provide a framework for understanding how lysosomal biogenesis has been maintained and adapted over a hundreds of million years of eukaryotic evolution.

## Material and Methods

### Phylogenomics and informatics

Pan-eukaryotic searches for all the surveyed genes were conducted using publicly available genome, transcriptome and protein datasets from The Comparative Set (TCS) of 196 species on EukProt v3 (Richter et al., 2022).Homology searches were conducted using tBLASTn, BLASTp and HMMer analysis into the TCS datasets using AMOEBAE (Analysis of Molecular Evolution with Batch Entry) pipeline designed for comprehensive comparative genomic survey (Barlow et al., 2023) supplemented with reciprocal searches against human database to confirm orthology (e-value ≤ 0.05) (Table S2-S3). Deep-dive comparative genomics (Table S3) was conducted using *T. vaginalis* and *A. ignava* sequences as queries for searches into all parasitic parabasalid databases freely available on NCBI. All the potential orthologues identified in the comparative genomic analyses were validated for the presence of conserved domains using InterProScan. Sequence alignments were conducted using MAFFT v7.505 (Katoh et al., 2019). Alignments were trimmed using Block Mapping and Gathering with Entropy (BMGE) using BLOSUM62 matrix and gap rate cut off (-h 0.8) to retain informative sites (Criscuolo & Gribaldo, 2010). IQTree v2.0 was used to conduct ultrafast bootstrapping (-B −1000) and a robust non-parametric bootstrapping with the calculation of Bayesian posterior probability values (-b 1000 -alrt 1000 -abayes) with best fit model predicted using ModelFinder (Minh et al., 2020).

Co-occurrence tests were conducted with custom R-script using Jaccard similarity index supplemented with Fisher’s exact test to compute similarity and diversity of the sample sets (Table S10). Jaccard indices were calculated using the mathematical formula J(A,B)= |A ∩ B| / |A ∪ B|, where A presents the set of species with protein A, B represents the set of species with protein B, |A ∩ B| represents number of species showing presence of both the proteins and |A ∪ B| represent number of species showing presence of either protein. Co-occurrence values (J) were calculated in a 13×13 matrix where J=0 represents proteins A and B never found together and J=1 represents both proteins always found together.

MRH assessment for M6P binding specificity was conducted using pair-wise sequence similarity analysis with manual detection of QREY tetrad and conserved cysteine residues (Figure S3B, Table S11). MRH domains with the highest pairwise similarity to an active human MRH (MRH3, 5, 9, 15 and CD-MPR) within a TvMPR, complete QREY score (100), and ≥6 cysteine residues represent high confidence in putative M6P binding ability. Alignment was graphically generated to highlight essential residues using Espript 3 (Gouet et al., 1999). Structural predictions and modelling were done using AlphaFold 3 (Abramson et al., 2024), while structural alignments were graphically generated on PyMOL version 3.0, LLC and mathematically supported by TM-score generated by TM-align (Zhang, 2005). Crystal structures for human MRH domains were obtained from protein data bank (PDB). All analyses were conducted on R-studio and python with self-made scripts using appropriate R-packages and Python packages provided in the open-sourced project on GitHub (https://github.com/abhishekpshinde1994/M6P-signaling-system-analyses.git).

Clustering of 329 MPR proteins, including CI-MPR and CD-MPR paneukaryotic homologs and parabasalid MPRs identified through comparative genomic surveys, was performed based on the all against-all pairwise sequence similarity using the CLANS (Frickey & Lupas, 2004). CLANS: a Java application for visualizing protein families based on pairwise similarity, is a web utility in the MPI Bioinformatics Toolkit (https://toolkit.tuebingen.mpg.de/tools/clans). The analysis was conducted using the default parameters (BLOSUM62 matrix and e-value = 1 × 10⁻⁴) of BLASTp. The CLANS graphical user interface (GUI), implemented in the Java Runtime Environment, was used for visualization with the default settings and clustering was allowed to proceed until overall clustering pattern appeared stable.

### Cell cultivation

*T. vaginalis* strain TV17-48 were cultured in Tryptone Yeast extract Maltose (TYM) medium at pH 6.2 supplemented with 10% inactivated horse serum at 37°C. Transfected cells were selectively grown using 200 µg/mL of Geneticin 418 corresponding to plasmid containing gene tagged with HA tag and 40 µg/mL Puromycin corresponding to plasmid containing gene tagged with V5 tag. *T. vaginalis* population in logarithmic phase of growth was used in all the experiments.

### Expression vector construction and transfection

All genes were PCR amplified from genomic *T. vaginalis* TV17-48 DNA (See Table S12 for primers). TvMPR3 (TVAG_498650), TvMPR7 (TVAG_255730), TvGNPT (TVAG_166090) and TvCath D (TVAG_336300) were cloned in vector pTagVag-HA-Neo that allow their expression in *T. vaginalis* with 2xhemagglutinin (HA) (Hrdy et al., 2004). TvMPR3 and TvMPR7 (TVAG_255730) was also cloned in vector pTagVag-V5-Puro and expressed with V5 tag (Stáfková et al., 2018). VPS26A/B were previously prepared with 2xHA tags and are repurposed here (Shinde et al., 2024). All genes were expressed under the control of their respective native promoter 300bp upstream of the gene coding sequence. The expression vectors were transfected into *T. vaginalis* TV17-48 strain cells using the Human T-cell Nucleofector kit (Lonza) according to (Zimmann et al., 2022).

### Generation of antibodies

Antigenic fragment of the MPR1 (TVAG_177320) protein sequence was selected based on most hydrophilicity and unique region showing least homology with other *T. vaginalis* MPR paralogs (fragment sequence in Table S12). The fragment was amplified and cloned into pet42B vector (Novagen) and expressed with 6xHis tag in *Escherichia coli* BL21 strain. The antigen was purified using Ni-NTA (Qiagen) affinity chromatography under denaturing conditions according to manufacturer protocol and used for development of rat polyclonal antibody (Biocev, Czech Republic). Similarly, protein disulfide isomerase (PDI; TVAG_076650) was expressed in *E. coli*, affinity purified and used for mouse polyclonal antibody development.

### Immunofluorescence microscopy and imaging

*T. vaginalis* TV17-48 wild-type (WT) and transfected strains were used for the immunofluorescence microscopy. *T. vaginalis* cells were harvested, washed with PBS and fixed with 2% formaldehyde as described previously (Woessner & Dawson, 2012).Cells were attached to coverslips coated with poly L-lysine followed by permeabilising with 0.2% Triton X-100. The cells were blocked with 1% BSA and incubated overnight with corresponding primary antibodies with appropriate dilutions: rabbit polyclonal anti-HA antibody (Exbio, 1:1000), mouse monoclonal anti-V5 antibody (Abcam, 1:1000), rat polyclonal anti-MPR1 antibody (1:250) and mouse polyclonal anti-PDI antibody (1:500). The tagged proteins were visualised with respective secondary antibodies conjugated with Alexa Fluor-488 and −594 (Life Technologies, 1:1000) for the co-localisation analysis. The nucleus was stained with 4′,6-diamidino-2-phenylindole (DAPI) in Vectashiled mounting medium (Vector laboratories). The cells were visualised using a Leica TCS SP8 confocal laser scanning microscope with a 63x oil immersion objective. For expansion microscopy, *T. vaginalis* cells were fixed with 1% formaldehyde, and expansion microscopy samples were prepared as described (Gorilak et al., 2021). Cell structures were labelled with Atto 594 NHS ester (ATTO-TEC, Siegen, Germany). Prepared samples were images using a Nikon CSU-W1 spinning disk confocal microscope (Nikon Corporation, Amstelveen, Netherlands) and images were processed as above. Z-stack images were captured and deconvolved using Huygens Professional v19.10, processing and analysis of cell images was done using Imaris 9.7.2 package and ImageJ/Fiji. Voxel based co-localisation analysis was performed using ImarisColoc to calculate Manders co-efficients and Person correlation co-efficient.

### Co-immunoprecipitation (coIP) of M6P signalling system proteins

Co-IP of TvMPR3, TvMPR7 and TvGNPT was performed as described previously using anti-HA antibody conjugated dynabeads (Exbio) (Kučerová et al., 2023; Makki et al., 2019) with and without cross-linking using WT and transfected strains. The experiments were conducted in biological triplicates for each bait protein. The bait-complex bound beads were processed for proteomic analysis described below.

### Label-free quantitative (LFQ) Mass spectrometry analysis

Peptides bound with each bait protein in the co-IP experiments were eluted and digested witth trypsin and purified as described previously (Masuda et al., 2008). Purified peptides were separated using nano-scale liquid chromatography (UltiMate 3000 RSLC, Thermo Scientific) using reverse-phase column (EASY-Spray column, 50 cm × 75 μm ID, PepMap C18, 2 μm particles, 100 Å pore size) and coupled with Orbitrap Fusion Tribrid mass spectrometer (Thermo Scientific), raw data was processed using MaxQuant software (version 1.6.3.4). The searches of the raw data were conducted with the precursor ion mass tolerance in the initial search was 20ppm, tolerance in the main search was 4.5ppm and the fragment ion mass tolerance was 0.5 daltons. Gene identification and annotations were obtained from UniProt and TrichDB databases. Quantifications were performed with label-free algorithm and data was presented as volcano plots with −10log-adjusted P-values of a false discovery rate (FDR) corrected t-test to be plotted against 2log fold-change of relative label-free quantification intensity. Significant enrichment was shown using cut-off with 2log fold-change (FC)>1 and P-value threshold of 0.05. Significantly enriched proteins with the bait protein are labelled as red dots, proteins enriched in the control co-IP are labelled as blue dots and insignificant proteins represented by grey dots. R-Script was written and used on R-studio to generate the volcano plots. Overlapping analysis and interactome network was built using R studio with Venn, ggvenn, ggplot2 and ggraph packages. R-scripts for both analyses can be found in the above mentioned open-sourced project on GitHub. The raw data has been deposited to the ProteomeXchange Consortium via the PRIDE with the dataset identifier PXD082344.

### Glycomics

N-glycans were released by PNGase F from Chinese hamster ovary (CHO) cells (ATCC CCL-61) and protein extracts of *T. vaginalis* TV17-48 strain cells, permethylated and fractionated on a Waters Oasis-MAX SPE cartridge for MALDI-MS and LC-MS/MS analyses as described previously for sulfoglycomic analysis of negatively charged sulfated glycans (Hoshino et al, 2024). All MS and MS^2^ data were manually interpreted and assigned based on comparison with those of similarly processed M6P-carrying N-glycans from CHO cells. Fragmentation patterns were illustrated by cartoon drawings using the standard symbol nomenclature (Varki et al. 2015).

## Acknowledgements

We thank Michaela Marcinčikova and Michal Havelka for their technical assistance. The authors acknowledge the Imaging Methods Core Facility at BIOCEV, Charles University, Czech Republic, a member of Euro-BioImaging ERIC and Czech BioImaging national infrastructure supported by MEYS LM2023050 grant – for their support and assistance in this work. The proteomic analyses were performed at the OMICS Mass Spectrometry Core Facility at BIOCEV, Faculty of Science, Charles University, we thank Jana Jasprova and Pavel Talacko for their expert assistance with proteomics. We also thank the Academia Sinica Common Mass Spectrometry Facilities for Proteomics and Protein Modification Analysis (grant AS-CFII-111-209) for MS data acquisition.

---

**Extended data Figure 1:**
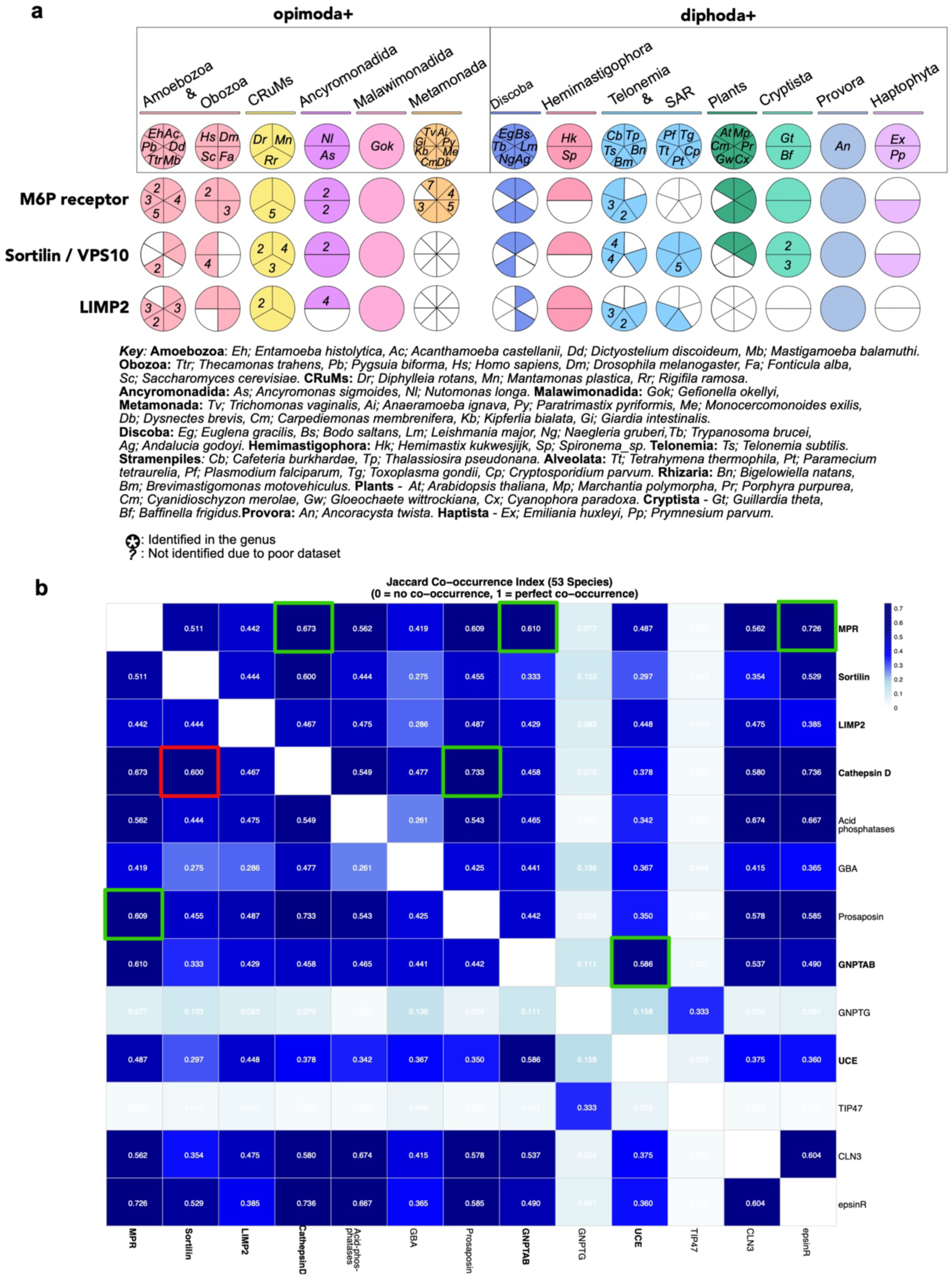
a) Coulson plot extended from Figure 1a demonstrating comparative presence of M6P independent receptors sortilin and LIMP2 with MPRs. Sections filled in colour demonstrate positive presence of a single protein homolog, empty sections demonstrate absence of protein homologs, numbers represent count of identified orthologues duplicated in the species. Abbreviated binomial nomenclature of each organism is de-abbreviated in the key. b) Jaccard index and co-occurrence test. Jaccard index values numerically shown indicating the possibilities of protein-protein co-occurrence across the 53 analysed species. Zero indicates no co-occurrence while 1 represents complete co-occurrence. Complete mathematical calculations are provided in Extended data Table S10. Significantly co-occurring relations are highlighted in green for M6P dependent pathway and in red for M6P independent pathway.

**Extended data Figure 2:**
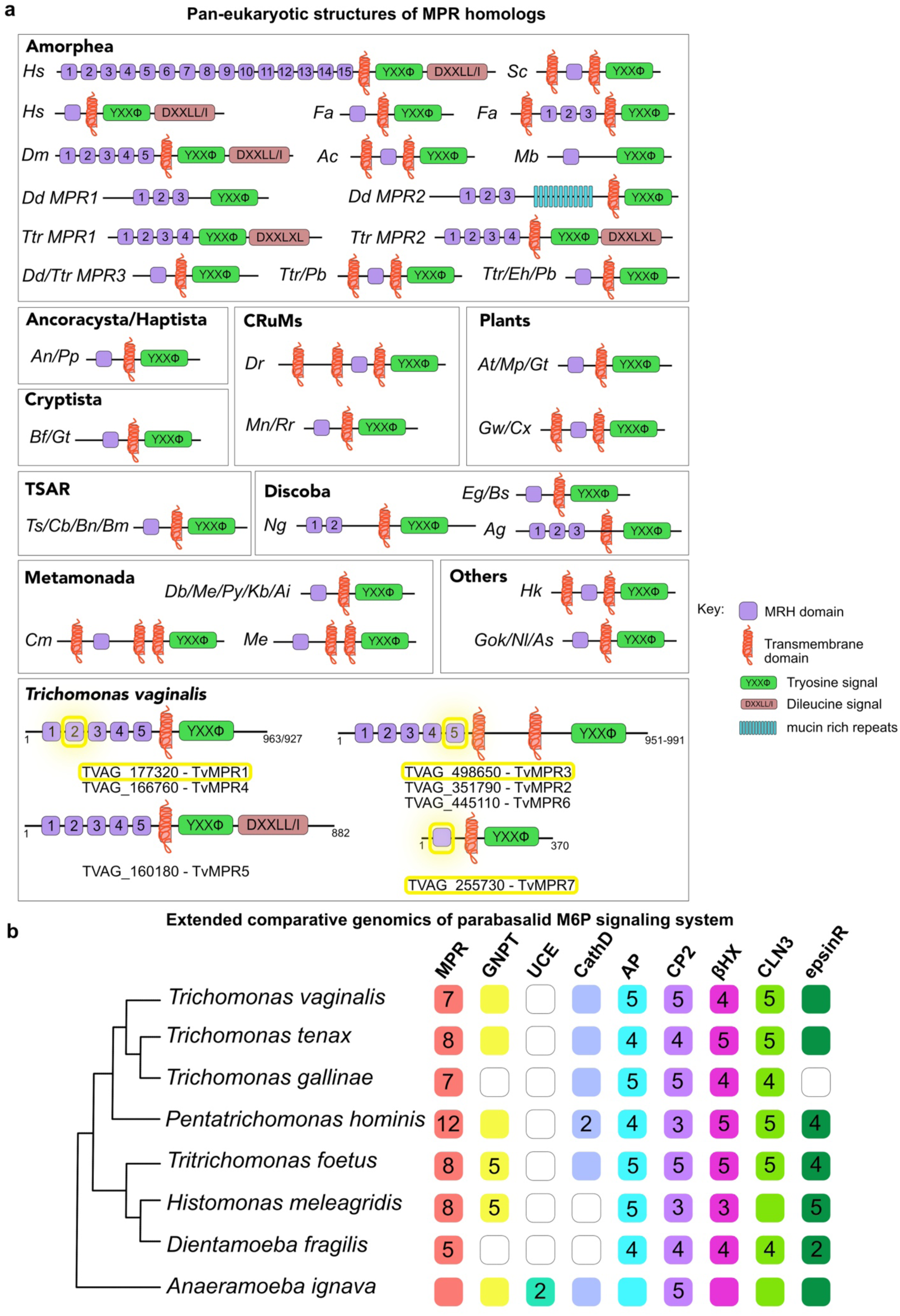
a) Domain topologies of MPR homologs identified across 53 pan-eukaryotic species classified based on their eukaryotic supergroup. Key is provided to describe each structural element. b) Extension to Figure 1b; comparative genomics of parabasalid M6P signalling system (refer to Figure 1b figure legend for description).

**Extended data Figure 3:**
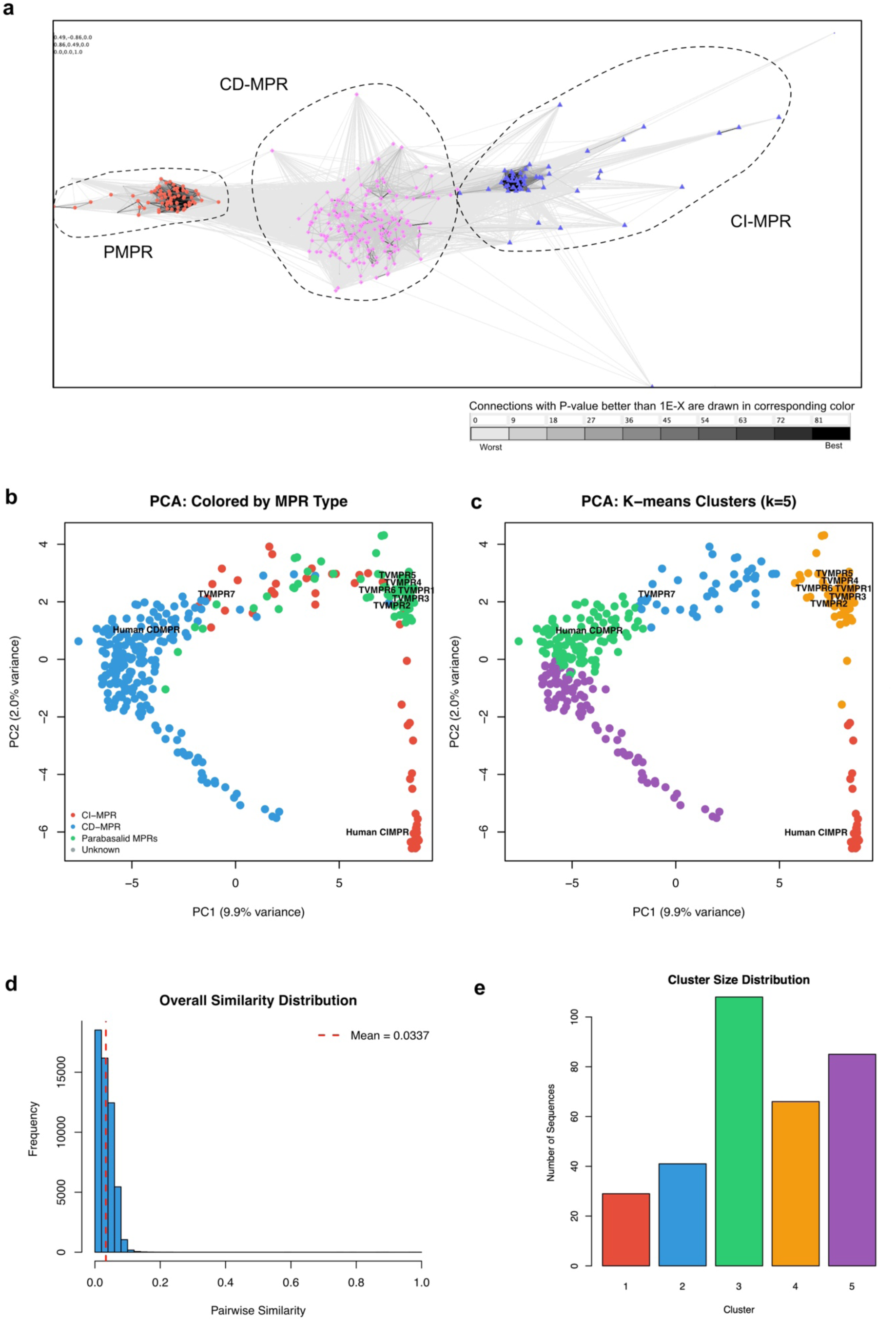
Pan-eukaryotic conservation across TCS species of MPRs. a) CLANS sequence similarity analysis. PMPR cluster with red nodes represent the proteins identified in parasitic parabasalids, CD-MPR cluster with pink nodes represents the CD-MPR homologs identified in all the eukaryotic species, CI-MPR cluster with blue nodes represents the CI-MPR homologs identified in all the eukaryotic species. b) Principal component analysis of MPRs categorised by MPR clustering (CI-MPR, CD-MPR, PMPR). c) PCA with k-means clustering analysis generating five pan-eukaryotic categories based on pairwise sequence similarity analysis. d) Pairwise similarity values versus frequency of occurrence plot demonstrating mean similarity of 0.0337 (3.37%). **e)** k-means cluster distribution with bar plot demonstrating number of sequences each cluster comparatively presents.

**Extended data Figure 4:**
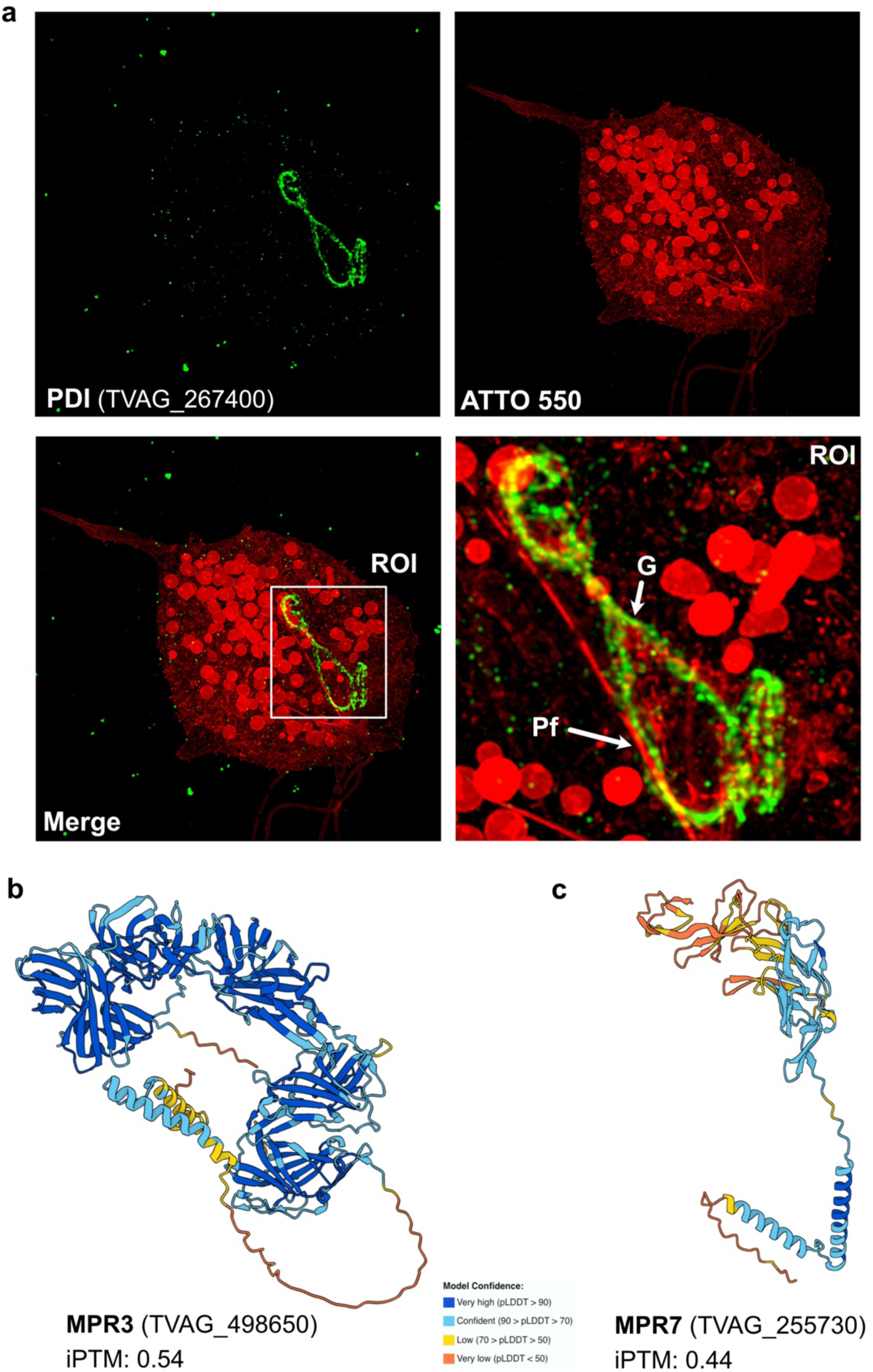
a) Expansion microscopy of Golgi marker; PDI. Green channel represents the labelling for α-PDI antibody marking the Golgi compartment in *T. vaginalis.* Red channel presents the staining of the cell structure with ATTO 550 NHS-ester. Merge channel presents overlapping signal with ROI zoomed in to demonstrate the proximity of Golgi (G) labelled with α-PDI with the parabasal filaments (Pf). b) Alphafold 3 prediction of TvMPR3 and TvMPR7, iPTM scores of overall structural confidences are provided in each panel, with pLDDT model confidence embedded graphically in the structure and presented as a key for both structures.

**Extended data Figure 5:**
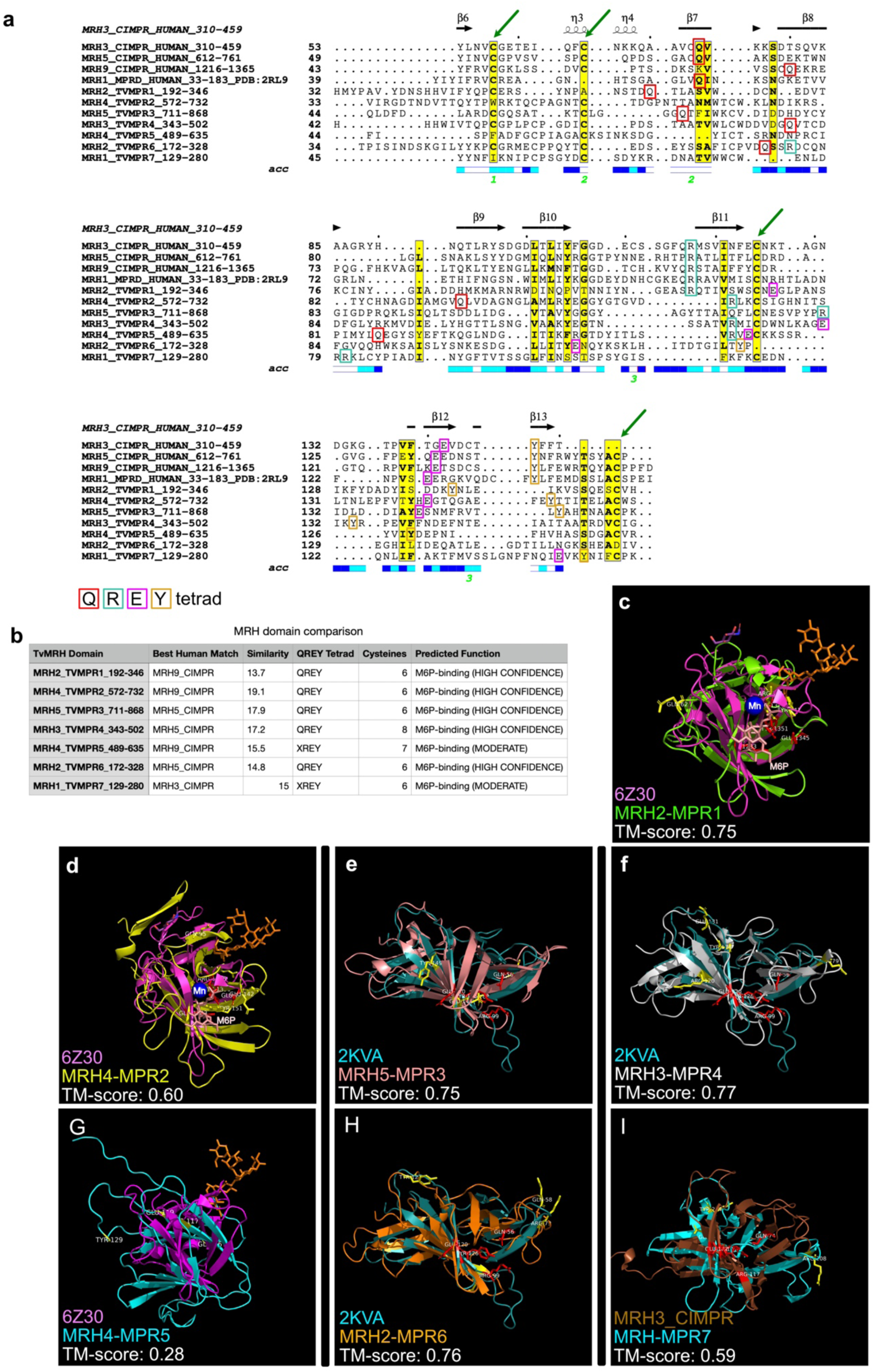
M6P binding predictions for TvMPR MRH domains a) Espript alignment of the highest scoring (on pairwise similarity matrix) MRH domains from each TvMPR paralog with functional human CI-MPR and CD-MPR MRH domains, with secondary structure annotation from CI-MPR MRH3 domain. Green arrows depict positions with conserved cysteine residues, Glutamine (Q), Arginine (R), Glutamic acid (E) and Tyrosine (Y) tetrads for each MRH domain in the alignment are annotated with colour squares (refer to key). b) TvMRHdomain confidence prediction for M6P binding, confident MRH domain comparison analysis values of each TvMPR (elaborated in Table S11). c-i) 3D structural alignment of highest scoring MRH domain from each TvMPR paralog with crystal structures of human MRH domains, alignment confidence is mathematically provided with the TM-align score in each panel, visual alignment is done on pyMOL.

**Extended data Figure 6:**
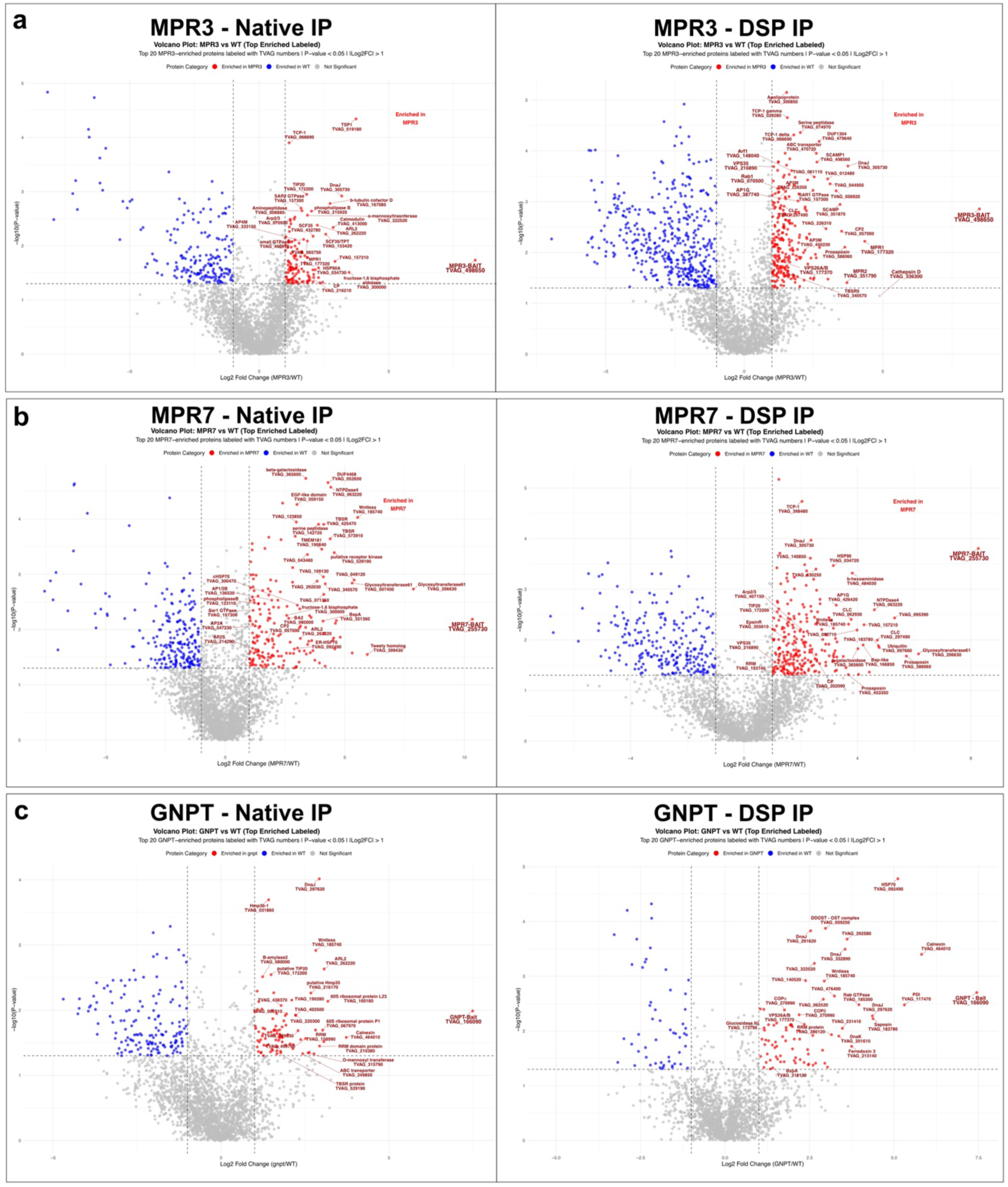
Volcano plots of analysed co-IP datasets. a) Volcano plots of TvMPR3 co-IP experiments under native and cross-linking (DSP) conditions. b) Volcano plots of TvMPR7 co-IP experiments under native and cross-linking (DSP) conditions. c) Volcano plots of TvGNPT co-IP experiments under native and cross-linking (DSP) conditions. Data in red represents significantly enriched proteins with the 2xHA tagged MPR3as bait, data in blue represents proteins enriched in the absence of bait, X-axis represents Log2 Fold change (TvMPR3 vs Wild type), Y-axis represents -log10 (p-value) for significance of enrichment within all three experimental replicates.

**Extended data Figure 7:**
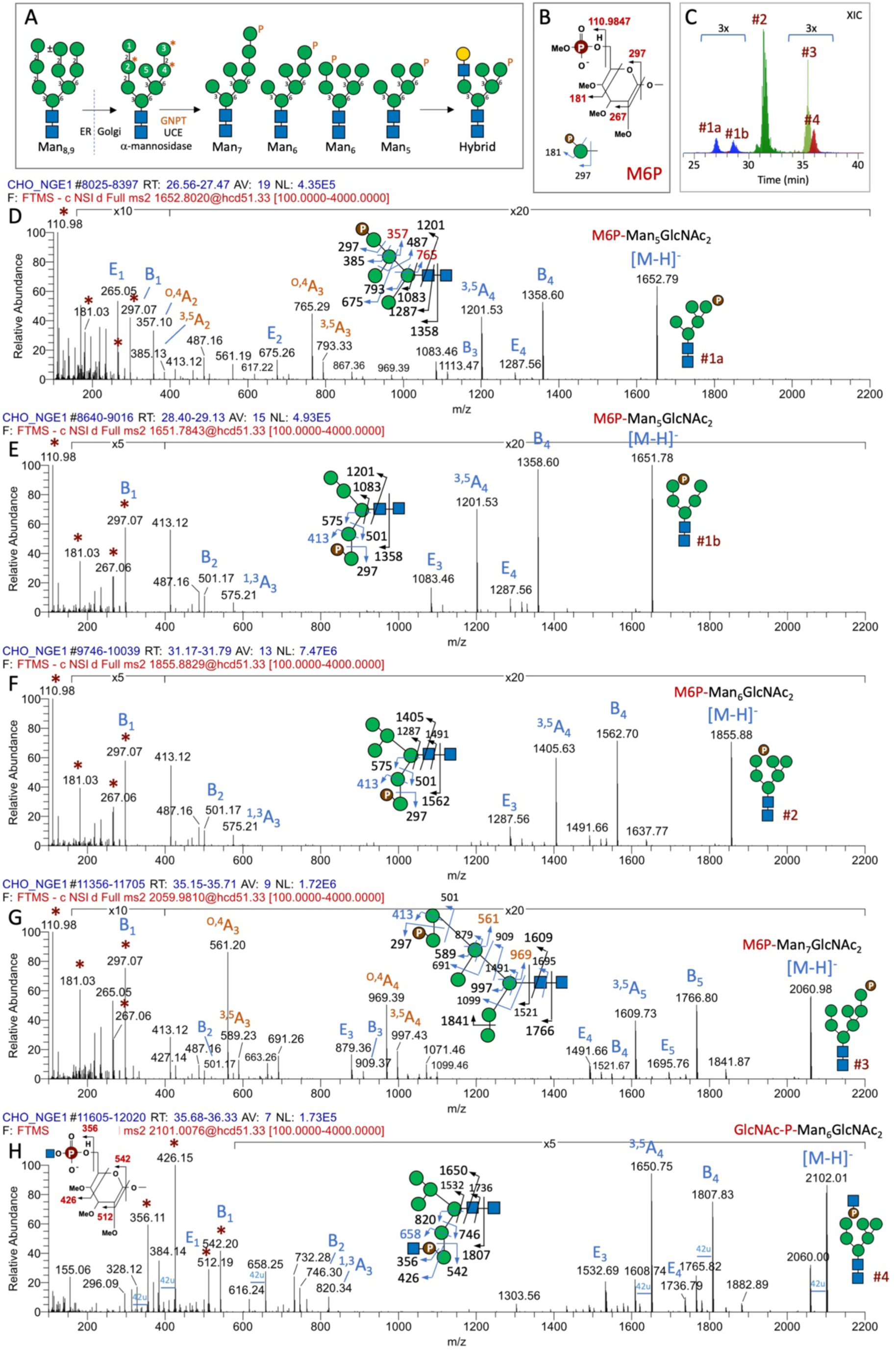
Identification of M6P-carrying N-glycans from CHO cells. In mammals, the 5 Man residues that can be GlcNAc-P-phosphorylated by GNPT are numbered 1-5, and the preferred sites were reported to be on residues 2,3 and 4 (Varki & Kornfeld, 1980). Subsequent removal of the GlcNAc by UCE and/or the non-M6P-capped Man by α2-mannosidase would generate the expected phosphorylated Man_5-7_high-mannose N-glycans as illustrated in the schematic drawing (panel A). Further removal of exposed, non-M6P-modified α3- and α6-Manresidues on the 6-arm of the trimannosyl core would normally require the addition first of a GlcNAc on the 3-arm by MGAT1. LacNAc extension can then be initiated on the 3-arm to generate the hybrid-type structures. For glycomic analysis, N-glycans released from total cellular protein extracts were reduced, permethylated, and the negatively charged permethylated M6P-carrying N-glycans eluted from an Oasis-MAX mixed-mode SPE were analyzed by LC-MS/MS in negative mode. The diagnostic ions at m/z 110.9847 (detected at 1-2 ppm mass accuracy on an Orbitrap), along with m/z 181, 267, 297 can be used to filter for terminal M6P-carrying glycans (panel B). The non-reducing terminal A, B, and E ions allow efficient sequencing of the underlying oligomannose structure, with the ring cleavage ^0,4^A ions critically defining C6 substitution(s) on the 6-arm, whereas the absence of it is indicative of M6P on the (±Manα1-2)Manα1-2Manα1-3-arm. Since only the phosphorylated, and hence negatively charged ions would be detected in negative mode MS^2^, the glycosyl residue where the cleavage occurs was numbered starting from the non-reducing terminal M6P-carrying Hex residue and denoted by the subscript following the ion type. Typically, the M6P-N-glycans that can be recovered and identified in the fraction containing singly negatively charged permethylated glycans by this glycomic approach (Yu et al., 2013) are the Man_6-8_GlcNAc_2_ structures, often dominated by the Man_6_ species carrying either a terminal M6P or GlcNAc-P. The latter would afford the diagnostic ions at m/z 356, 426, 512, and 542 instead, corresponding to a shift of 245 u due to the additional GlcNAc. The MS^2^ fragmentation patterns for the identified M6P-carrying N-glycan structures from CHO cells (peaks #1-4 in the overlaid extracted ion chromatograms shown in panel C) are illustrated with schematic drawings and peak annotations in their respective MS^2^ spectra (panels D-H). There are likely additional isomeric forms of the M6P-Man_5_GlcNAc_2_ coeluting with peak #1a, but the major form is defined by the two ^0,4^A ions as illustrated in (D). Notably, the terminal (M6P)Man-2Man-moiety produced a characteristic fragment ion at m/z 413 (see MS^2^ spectra for peaks #1b, 2, 3) that was shifted to m/z 658 for (GlcNAc-P)Man-2Man-(peak #4). This is not a typical ring-cleavage A ion (exact structure unclear) but is nonetheless highly sequence-informative. Additional doubly negatively charged Man_6,7_high-mannose N-glycan structures carrying two M6P or GlcNAcP modifications can be found and identified by the same manner in the later eluting fractions from the OasisMax SPE (not shown).

**Extended data Figure 8:**
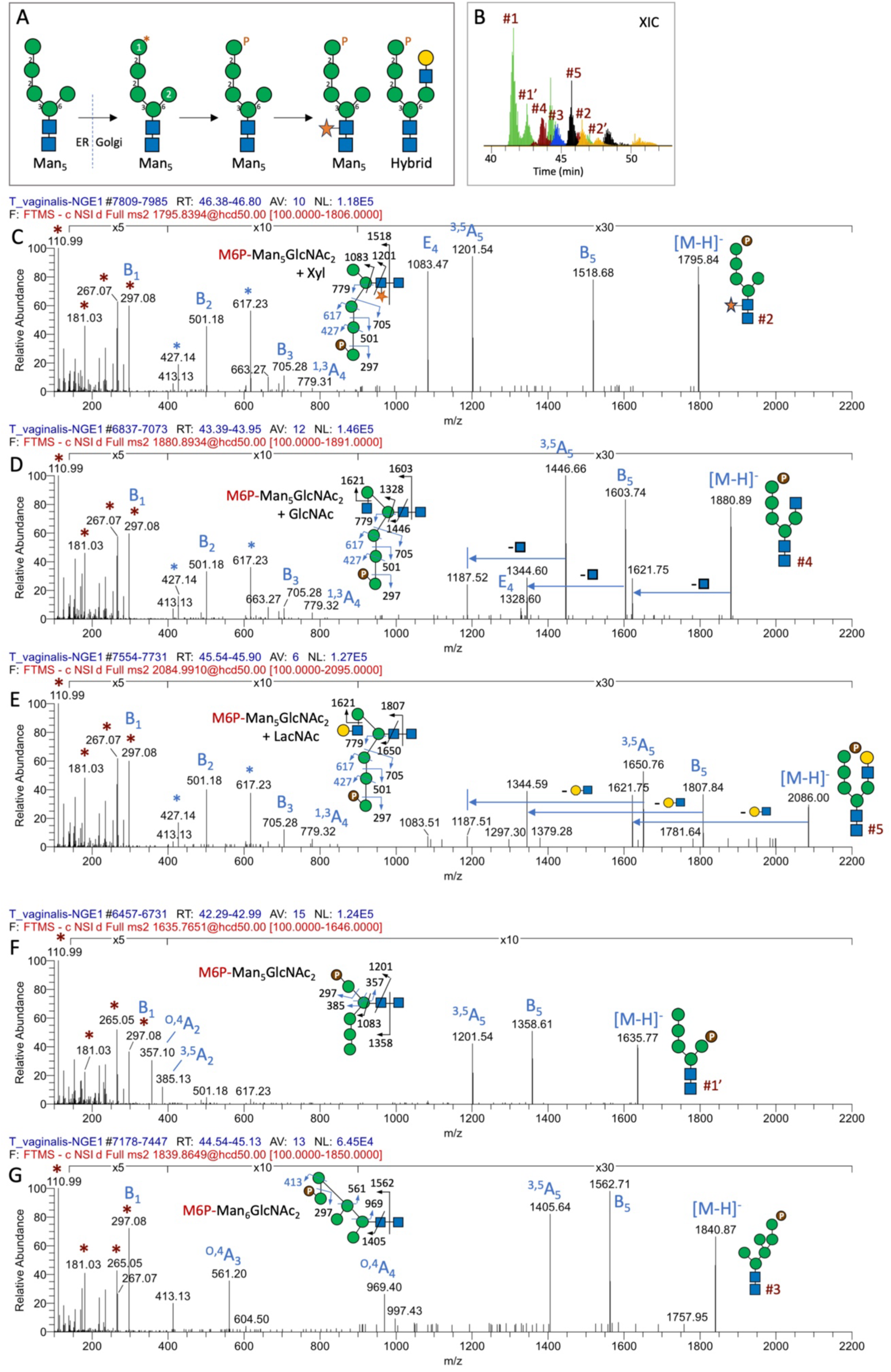
M6P-carrying N-glycans from *Trichomonas vaginalis*. In contrast to the mammalian system (Extended data Figure 7), the dolichol-linked precursor synthesized in *T. vaginalis* is not further extended on the 6-arm due to the absence of mannosyltransferases in ER lumen (Samuelson et al., 2005). The high mannose N-glycan made is therefore a unique Man_5_GlcNAc_2_structure distinct from that typically found in mammals, which can be extended on the 6-arm by GlcNAc and LacNAc, or further core xylosylated (Paschinger et al., 2012). Each of these Man_5_GlcNAc_2_-based structures could carry a M6P at the terminal Man (panel A), as identified in this work by negative mode LC-MS/MS analysis of the permethylated negatively charged N-glycans. Since the released N-glycans were not reduced, each species yielded two anomeric peaks on LC, as shown by the overlaid extracted ion chromatogram (panel B). Peaks #1 and 4 were each further resolved into a major and a trailing peak (peaks #1’ and 4’). The MS^2^ spectrum for the most abundant M6P-Man_5_GlcNAc_2_ component (peak #1) is shown in Figure 6B, while those of peaks #2-5 are shown here (panels C-G). The diagnostic MS^2^ ions at m/z 110.9847, 181, 267, and 297 afforded by all peaks confirm that they all carry a terminal P-6Hex. Instead of the characteristic fragment ion at m/z 413 (m/z 297+116 u) produced by structures carrying a terminal (M6P)Man-2Man-moiety as found in the CHO cells (Extended Data Fig. 7), the linear (M6P)Man-2Man-2Man-moiety extending from the 3-arm yielded unique fragment ions at m/z 413/427 and 617 (m/z 505+116 u). It is similarly unclear how these ions were formed but their common presence alongside the B_1_, B_2_, B_3_, and ^1,3^A_4_ions defines an identical Man_5_GlcNAc_2_ core structure substituted with an additional Xyl (peak #2) or extended at the 6-arm by a GlcNAc (peak #3) or LacNAc (peak #4). In contrast, the absence of these ions but presence of the ^0,4^A_2_ and ^3,5^A_2_ ions at m/z 357 and 385 indicate that the same Man_5_GlcNAc_2_ structure could alternatively carry the M6P on the 6-arm Man (peak #1’). Intriguingly, a Man_6_GlcNAc_2_ structure (peak #5) was found to have a 6-arm extension similar to that identified in CHO cells (peak #3, Extended Data Fig. 7G), but inconsistent with the currently understood biosynthesis of dolichol-linked precursor in *T. vaginalis*.

**Extended data Figure 9:**
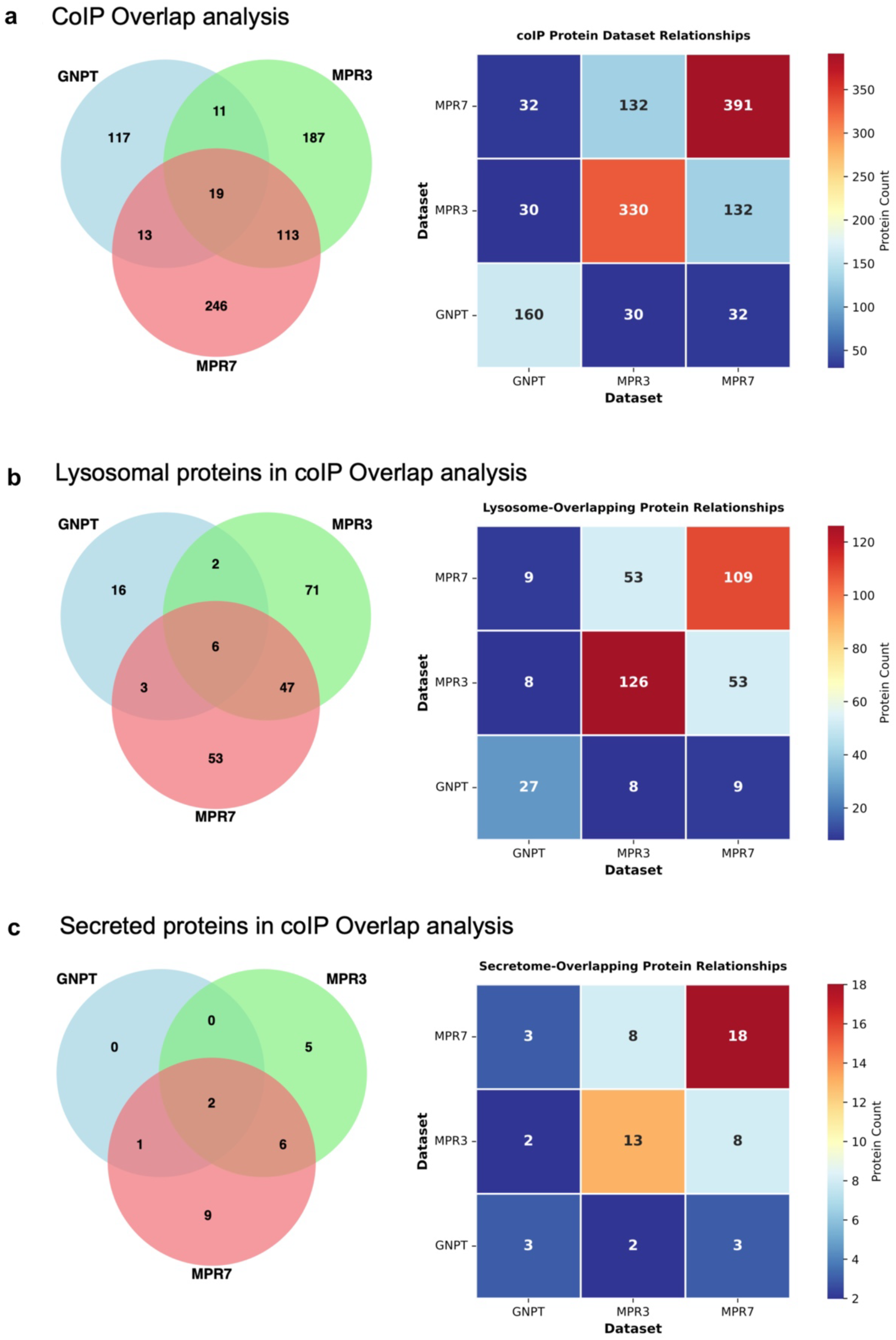
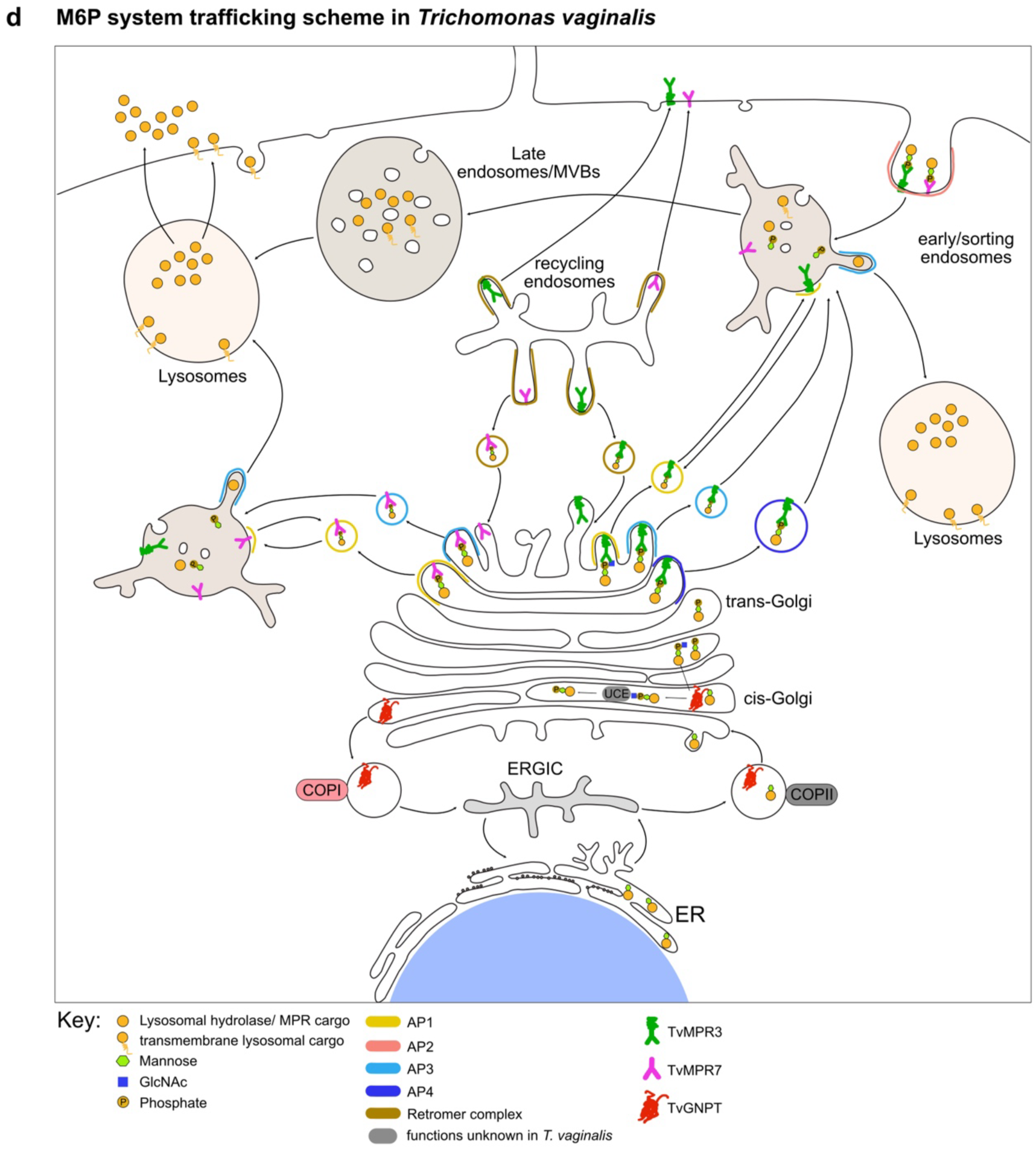
Quantification of phagolysosomal and dynamically secreted proteins and schematic of M6P signalling pathway. a) Overlap of significantly enriched protein identifications with all three baits as presented by a Venn diagram and relationships of significantly co-occurring within each bait pair. b) Overlap of significantly enriched phagolysosomal proteins in the interactome presented by a Venn diagram and their relationships within bait pairs. c) Overlap of significantly enriched secreted proteins in the interactome presented by a Venn diagram and their relationships within bait pairs. d) Schematic representation of M6P signalling generation and trafficking via distinct TvMPRs in *T. vaginalis*.

